# Transcriptomic and chromatin accessibility profiling unveils new regulators of heat hormesis in *Caenorhabditis elegans*

**DOI:** 10.1101/2025.03.11.642714

**Authors:** Hsin-Yun Chang, Sarah E. McMurry, Sicheng Ma, Charles L. Heinke, Christian A. Mansour, Sophia Marie T. Schwab, Charles G. Danko, Siu Sylvia Lee

## Abstract

Heat hormesis describes the beneficial adaptations resulting from transient exposure to mild heat stress, which enhances stress resilience and promotes healthy aging. While heat hormesis is widely observed, much remains to be learned about its molecular basis. This study bridges a critical knowledge gap through a comprehensive multiomic analysis, providing key insights into the transcriptomic and chromatin accessibility landscapes throughout a heat hormesis regimen in *C. elegans*. We uncover highly dynamic, dose-dependent molecular responses to heat stress and reveal that while most initial molecular changes induced by mild stress revert to baseline, key differences emerge in response to subsequent heat shock challenge that likely contribute to physiological benefits. We further demonstrate that heat hormesis extends lifespan specifically in wild-type animals, but not in germlineless mutants, likely due to transient disruption of germline activities during mild heat exposure, which appears sufficient to trigger pro-longevity mechanisms. This finding points to tissue-specific responses in mediating the physiological outcomes of heat hormesis. Importantly, we identify several highly conserved regulators of heat hormesis that likely orchestrate gene expression to enhance stress resilience. Among these regulators, some (MARS-1/MARS1, SNPC-4/SNAPc, FOS-1/c-Fos) are broadly required for heat hormesis-induced benefits, whereas others (ELT-2/GATA4, DPY-27/SMC4) are uniquely important in specific genetic backgrounds. This study advances our understanding of stress resilience mechanisms, points to multiple new avenues for future investigations, and provides a molecular framework for promoting healthy aging through strategic mid-life stress management.

## Introduction

Transient exposure to mild stress is well known to activate adaptive mechanisms that confer long-term protective effects and promote health, a phenomenon known as hormesis (1–4). Specifically,’heat hormesis’ describes the beneficial outcomes triggered by mild thermal stress, documented across species from yeast to humans (4–13). Sauna therapy, for instance, is increasingly recognized for its potential to improve health and mitigate age-related diseases through mechanisms attributed to heat hormesis (14,15). Despite extensive observational evidence of the beneficial effects of heat hormesis across diverse species and growing popularity in wellness practices, the molecular mechanisms underlying heat hormesis remain incompletely understood. *C. elegans* represents an ideal model for dissecting heat hormesis mechanisms due to its genetic tractability, well-characterized stress responses, and short lifespan.

In *C. elegans,* exposure to elevated temperatures at various life stages, including short exposures in young adulthood(4,6,16–19) or chronic exposure throughout development (20,21), consistently boosts thermal resistance and extends lifespan. Genetic analyses reveal a critical role of the transcription factor HSF-1 in mediating heat hormesis (17,19,22–25). HSF-1 is well-established to mount a robust transcriptional program in response to heat stress, which includes rapid induction of heat shock protein (hsp) genes.

Additional stress response transcription factors, such as DAF-16/FOXO, HIF-1, HLH-30/TFEB, have also been implicated in heat hormesis (19,20,22–27). Recent studies also highlight chromatin regulators, such as histone acetyltransferase CBP-1 and histone remodeler SWI/SNF, as participants in the long-term beneficial effects of heat hormesis (21,28).

Using transcriptomic profiling, a recent study identified distinct patterns of gene expression changes of *C. elegans* after a short exposure to heat stress (35°C for 1 hour) and 4 hours after recovery at normal growth temperature, including genes whose expression persists throughout the 4 hours of recovery, as well as genes that continue to be induced during recovery (29). Interestingly, some changes specific to the recovery period are regulated by the endoribonuclease ENDU-2, independent of HSF-1 (29). Another transcriptomic study comparing *C. elegans* raised at 15°C versus 25°C (as a model of chronic stress exposure) revealed differential gene expression profiles between the two populations (21), indicating *C. elegans* exhibits specific gene expression patterns in response to different temperatures.

Despite these advances, a longitudinal multiomic analysis that tracks changes across the entire heat hormesis regimen, including subsequent stress challenge, remains unexplored. Such data could provide crucial molecular insights that correlate with the improved physiological outcomes observed under hormesis conditions and identify possible regulators of the protective effects. This study fills this gap by presenting detailed transcriptomic and chromatin accessibility profiles at key timepoints throughout a heat hormesis regimen. We find that mild heat exposure induces extensive changes in RNA expression and chromatin accessibility that, although largely restored after a recovery period, leave distinct molecular imprints upon subsequent heat shock (HS). Our findings illustrate the dynamic molecular landscape during heat hormesis, providing concrete molecular evidence for dose-dependent responses and uncovering new candidate regulators with critical roles in diverse biological functions. Notably, multiomic analyses in wild-type (WT) and the germlineless mutant *glp-1(ts)* point to transient disruption of germline activities during mild heat stress exposure, which contributes to longevity extension, indicating tissue-specific responses to heat stress can have lasting effects on organismal physiology.

Furthermore, among the candidate regulators we uncovered, some show context-dependent regulation of heat hormesis, while others are broadly important. Our study offers valuable insights into the molecular basis for the development of stress resilience and may pave the way for promoting healthy aging through stress management.

## Results

### Priming induces thermal-resistance

To understand the molecular basis of heat hormesis, we adapted a regimen in which *C. elegans* that had been cultured at 20°C were exposed to 30°C for 6 hours during early adulthood (referred to as ‘priming’), allowed to recover at 20°C, and then challenged with heat shock (HS) at 35°C or 37°C. We applied this hormesis regimen to wild-type (WT) or *glp-1(e2144)* mutant (which become germlineless when grown at the non-permissive temperature of 25°C, hereafter referred to as *glp-1(ts)*) worms, and tested their thermal-resistance after HS.

We found that priming significantly improved thermal-resistance, based on motility scoring, in WT and *glp-1(ts)* mutant worms. The enhanced thermal-resistance persisted after 12-or 48-hours of recovery at 20°C (Fig. 1b), but the effect waned after a 96-hour recovery period. These results indicated that both WT and *glp-1(ts)* worms can retain a “memory” of the priming experience for an extended period of time, thereby exhibiting enhanced resistance to subsequent HS challenge. We tested varying priming durations and determined that a 6-hour priming showed substantially greater protective effect compared to shorter priming times (Fig. 1c). We additionally monitored survival after HS and observed that the primed worms lived substantially longer than their naive counterparts post HS (Fig. 1d-e; Supplementary Data 4).

**Figure 1:**
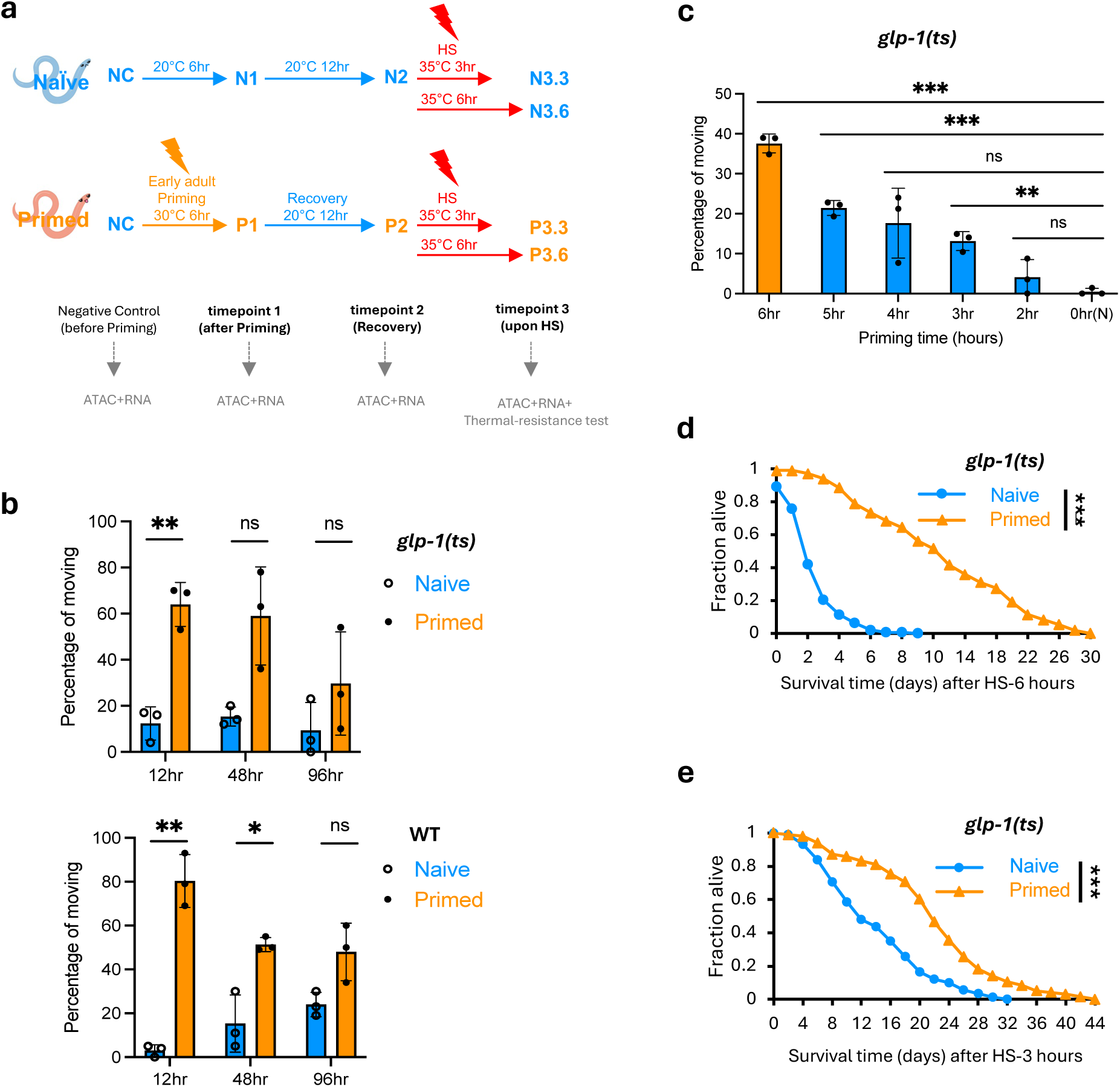
A heat hormesis regimen robustly enhances thermal-resistance. **(a)** Schematic of heat hormesis regimen and timepoints of multiomics studies: The primed group was exposed to 30°C for 6 hours (priming) during the early adult stage, followed by 12 hours of recovery at 20°C. The naive group was incubated at 20°C concurrently. Both groups were subjected to subsequent heat shock (HS) challenge at the indicated temperature and duration. Negative control (NC) timepoint was collected before priming. For both naive and primed groups, timepoint 1 (T1) was collected after priming (N1 and P1), timepoint 2 (T2) after recovery (N2 and P2), and timepoints 3 (T3) after 3 hours or 6 hours of HS (N3.3, P3.3, N3.6, P3.6). (**b)** Thermal-resistance of *glp-1(ts)* (*e2144* temperature-sensitive allele) (top) and WT (bottom), measured by the percentage of worms moving 15 h post-HS. The y-axis represents average results from three separate experiments, each with a different recovery period between priming and subsequent HS challenge. Recovery times and HS conditions varied due to the ease of experimental design. *glp-1(ts):*12 h recovery followed by 10 h of HS at 35°C; 48 h recovery followed by 5 h of HS at 37°C; 96 h of recovery followed by 5 h of HS at 37°C (N = 3); WT: 12 h recovery followed by 6 h HS at 35°C; 48 h recovery followed by 4 h HS at 37°C; 96 h recovery followed by 4 h HS at 37°C (N = 3). **(c)** Thermal-resistance of *glp-1(ts)* worms underwent various priming durations at 30°C, followed by 12 h recovery at 20°C, prior to a 10-hour HS at 35°C. Movement was scored 15 hours post-HS. Statistical analyses were conducted by comparing the indicated priming time to the 0-hour (Naive). (N = 3). (**d-e)** Survival of *glp-1(ts)* worms after HS 6 h (**d**) or 3 h (**e**) at 35°C, following a 12 h recovery from priming. The figure represents combined data from three independent experiments (N = 3). A 2-tailed unequal variances t-test was conducted to compare differences between naive and primed groups in (b-e). ** Indicates p < 0.01, *** Indicates p < 0.001. Details can be found in *source.data*.

### Transcriptomic and chromatin accessibility profiling in heat hormesis

To investigate the molecular basis of how transient exposure to mild heat stress, i.e., priming, confers resistance to a more intense heat shock (HS) later, we profiled mRNA expression and chromatin accessibility using RNA-seq and ATAC-seq, respectively, at key timepoints across our heat hormesis regimen. We hypothesized that RNA-seq and ATAC-seq data together would provide a more comprehensive view of the dynamics of gene expression regulation across the heat hormesis regimen. We initially conducted the multi-omic analyses using *glp-1(ts)* mutant, which lacks germline cells, thus enabling the assessment of molecular changes in somatic cells, and minimizing the possible confounding effects associated with extensive gene expression dynamics during early adulthood of reproductive worms. The timepoints we chose, including immediately after the 6-hour priming at 30°C (timepoint 1), after the 12-hour recovery at 20°C (timepoint 2), and after 3 or 6 hours of HS at 35°C (timepoint 3) (Fig. 1a), were guided by the dramatic phenotypic differences between primed and naive worms after HS (Fig. 1d-e).

We first examined RNA expression and chromatin accessibility changes in the naive group. We detected minor changes at timepoints 1 and 2 (N1 vs NC, N2 vs N1) (Supplementary Fig. 2a), likely reflecting developmental progression at normal culturing temperature. HS induced dramatic changes both in RNA expression and in chromatin accessibility (N3 vs N2) (Supplementary Fig. 2a; *source.data*). These changes showed significant overlap with previously reported heat stress-induced gene expression profiles (Supplementary Fig. 2e; *source.data*)(29,30), despite differences in experimental setup, supporting the validity of our results.

### Priming-responsive changes largely restore after recovery

We next focused on the primed group: Both RNA-seq and ATAC-seq revealed substantial changes immediately after priming (P1 vs NC), with 1,390 genes and 808 peaks significantly upregulated and 1,423 genes and 2,110 peaks downregulated, respectively (Fig. 2a–b, top; Supplementary Fig. 2b).

**Figure 2:**
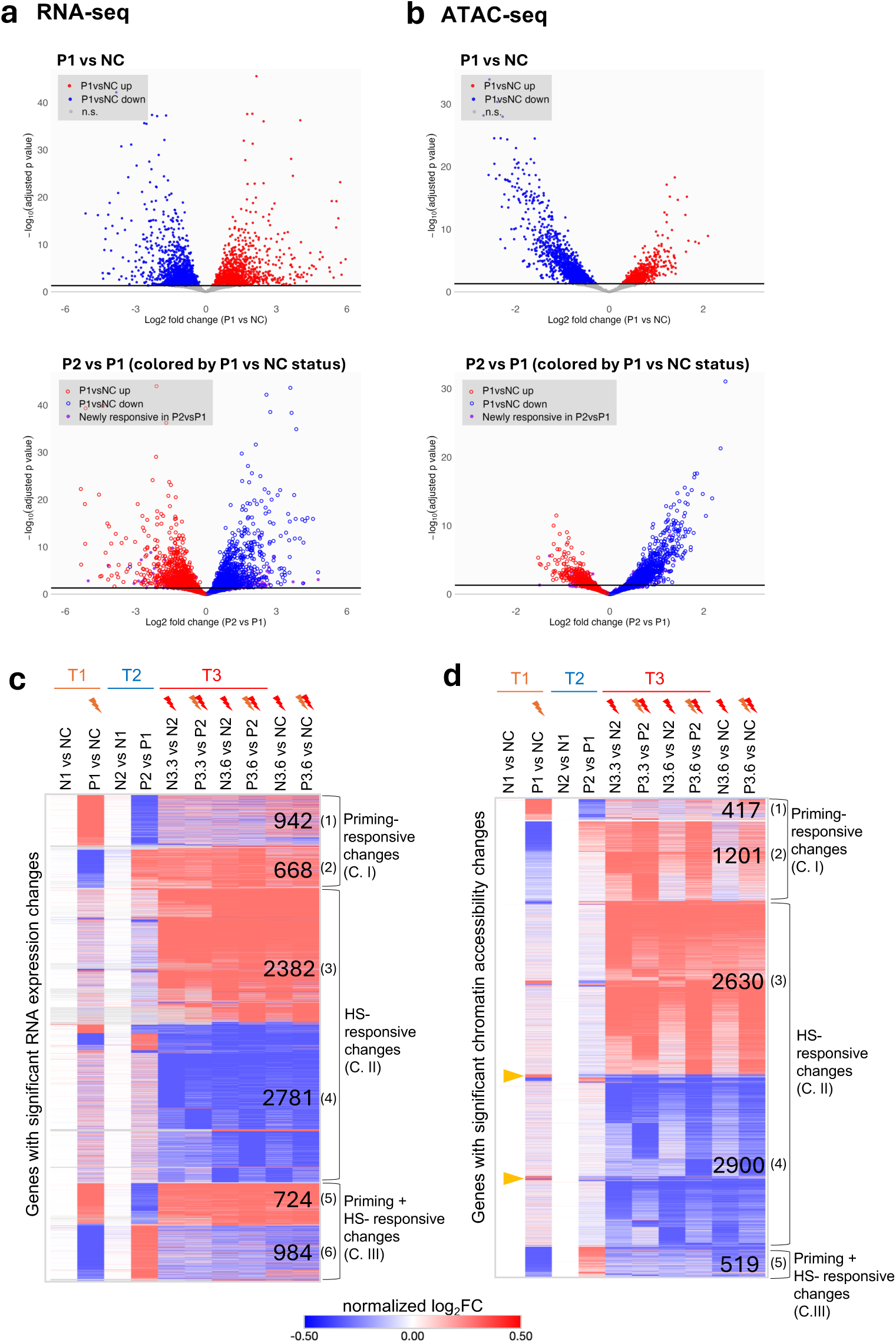
Priming-responsive changes largely restore after recovery and are distinct from Heat Shock-responsive changes in gene expression and chromatin accessibility. (a) RNA-seq and **(b)** ATAC-seq volcano plots. Top panels: Differential gene expression or chromatin accessibility after priming (P1 vs NC). Red and blue indicate significantly up-or downregulated genes/peaks, respectively (adjusted p < 0.05, threshold shown as black horizontal line). Bottom panels: Differential gene expression or chromatin accessibility after recovery (P2 vs P1), with genes/peaks colored by their P1 vs NC status. Red and blue open circles represent genes/peaks previously up-or downregulated at P1 vs NC, showing largely opposite regulation after recovery. Purple points indicate “newly responsive” genes/peaks that were not significantly changed in P1 vs NC but became differentially regulated in P2 vs P1. For RNA-seq **(a),** the y-axis was truncated for visualization (Max = 45). Complete gene and peak lists are provided in source.data. Heatmaps display genes with significant changes in RNA expression **(c)** or genes linked to the regions with significant change in chromatin accessibility **(d)** across the indicated comparisons, clustered using k-means in Morpheus. Panel **(c)** includes 8,481 genes, and panel **(d)** includes 10,837 genes linked to 17,943 chromatin regions (peaks). Colors represent normalized log2FC. Numbers indicate the size of each cluster (genes in **c**; genes with associated peaks in **d**). The clusters were rearranged to highlight shared patterns between transcriptomic and chromatin accessibility changes. Heatmaps are organized into three categories: Category I (C.I), Priming-responsive changes, including clusters 1–2 in **(c-d)**; Category II (C.II), HS–responsive changes, including clusters 3–4 in **(c-d)**; and Category III (C.III), Priming + HS-responsive changes, including clusters 5–6 in **(c)** and cluster 5 in **(d).** A small subset of gene-associated peak regions (yellow arrowheads in **d**) gained accessibility upon priming but not during HS, more closely resembling the pattern of RNA-seq cluster 1. Details of the gene lists in the heatmaps can be found in the Supplementary Data 1 & 2.

Strikingly, the vast majority of these priming-responsive changes reversed direction after a 12-hour recovery at 20°C (P2 vs P1) (Fig. 2a-b, bottom), indicating that the transcriptional and chromatin accessibility programs induced by priming are transient and reversible.

Upon HS at 35°C, we observed dramatic changes in RNA expression and chromatin accessibility in the primed group (Supplementary Fig. 2b), similar to that in the naive group as discussed above (Supplementary Fig. 2a). (Details of the gene and peak lists are in the *source.data*) We also compared the heat shock timepoint (T3.6) directly to baseline (NC), in addition to the timepoint after recovery. As expected, the two comparisons yielded nearly identical patterns (Fig. 2c-d), supporting our conclusion that most priming-induced changes are transient and return to basal levels by timepoint 2.

A comparison between the RNA-seq and ATAC-seq results indicated that there was a significant overlap, as well as distinction, between the genes associated with RNA expression or chromatin accessibility changes through the hormesis regimen. The data together supported our notion that the two genomic assays together provided a more comprehensive view of the gene regulatory changes associated with our heat hormesis regimen.

### Different levels of heat stress induce largely distinct RNA or chromatin accessibility changes

To further characterize the dynamic changes in RNA expression and chromatin accessibility across our hormesis regimen, we performed K-means clustering analysis of the genes associated with significant changes in RNA-seq or ATAC-seq across the three timepoints in either the naive or primed groups, and the results were visualized using heatmaps. The heatmap of ‘Genes with significant RNA expression changes’ includes the total number of genes identified from RNA-seq data (Fig. 2c, Supplementary Data 1), while the heatmap of ‘Genes with significant chromatin accessibility changes’ includes the genes associated with the peaks identified from ATAC-seq data (Fig. 2d, Supplementary Data 2). Interestingly, despite RNA-seq and ATAC-seq revealing substantially different genes associated with significant changes across the hormesis regimen (Supplementary Fig. 3c), the clustering analyses revealed a similar pattern of changes (Fig. 2c-d). We classified these shared patterns into three categories: Priming-responsive changes (C. I), HS-responsive changes (C. II), and Priming + HS-responsive changes (C. III) (Fig. 2c-d).

Priming-responsive changes (C.I) describe those exhibiting either upregulation or downregulation after priming (P1 vs NC) and showed the opposite trend following recovery (P2 vs P1), again supporting our earlier conclusion that most of the priming-induced RNA and chromatin accessibility changes are restored after recovery. Changes in this category were not robustly recapitulated after HS (timepoint 3) (Fig. 2c-d, Clusters (1) & (2)). The HS-responsive changes (C.II) displayed robust changes in RNA expression or chromatin accessibility upon HS (timepoint 3.3 or 3.6 vs timepoint 2), with minimal alterations observed after priming (timepoint 1) and following recovery (timepoint 2) (Fig. 2c-d, Clusters (3) & (4)).

The final category (C. III) includes genes that responded to both priming and HS, with most exhibiting consistent up-or down-regulation upon priming and HS (Fig. 2c-d, Clusters (5) & (6)). Interestingly, in this category, ATAC-seq data uncovered only genomic regions associated with significantly reduced chromatin accessibility upon priming and HS (Fig. 2d). Close inspection revealed that many regions in ATAC-seq Cluster 1 showed a strong accessibility increase after priming, but a weaker, yet still detectable, increase after HS (resembling the behavior of RNA-seq Cluster 5). We also observed a small set of regions that gained accessibility upon priming but not during HS (highlighted in Fig. 2d), which more closely resemble RNA-seq Cluster 1. However, the limited number of these regions prevented them from forming a distinct cluster under our k-means approach.

To understand the potential biological relevance of the various changes, we conducted Gene Ontology (GO) analyses of the various clusters. For Priming-responsive changes (C. I), we note that ‘ribosome-related’ and ‘stress response’ were the functional groups associated with both higher and lower expression and more and less chromatin accessibility, perhaps reflecting the dynamic nature of these classes of genes during the priming period (Supplementary Fig. 3a). Similarly for HS-responsive changes (C. II), ‘lipid metabolism’ and ‘small GTPase signaling’ were the functional groups represented by both more and less chromatin accessibility. Interestingly, ‘NHR transcription factors’ were associated with both downregulated RNA expression and less open chromatin (Supplementary Fig. 3b). For the changes that are shared between priming and HS (C.III, Priming + HS-responsive changes), ‘mRNA processing’, ‘heat stress response’, and ‘proteolysis proteasome’ were the top significantly enriched functional groups associated with genes with upregulated RNA expression (848). We note that canonical heat shock response genes are among this group, and their repeated induction likely contributes to the phenotypic protection conferred by heat hormesis. ‘Metabolism (lipid, short chain dehydrogenase)’ and ‘detoxification stress response’ were among the most significantly enriched GO terms for genes with downregulated RNA expression (980). Additionally, ‘metabolism (lipid and amino acid)’ was significantly enriched for genes with less open chromatin (567) (Supplementary Fig. 3c). The downregulation of lipid metabolism genes may reflect a compensatory response to changes in membrane fluidity at high temperature. (Details of the GO analysis can be found in Supplementary Data 3)

### Primed worms exhibit differential RNA expression and chromatin accessibility upon HS compared to naive worms

To uncover the molecular basis underlying the protective effects of primed worms, we assessed the differences in RNA expression and chromatin accessibility between primed and naive groups at the various timepoints. At timepoint 1, our finding was similar to what was discussed earlier, since naive worms did not show substantial changes, so changes between primed and naive were largely similar to those detected in primed worms comparing P1 vs. NC (Fig. 3a, Supplementary Fig. 2b; *source.data*). At timepoint 2, we detected only a handful of significant differences between primed and naive worms, which is consistent with our earlier conclusion that the majority of the priming-induced changes were restored after recovery (Fig. 3b; *source.data*). Nevertheless, we identified a small subset of genes that exhibited significantly persistent RNA expression change through recovery (30 among 1501 upregulated, 43 among 1432 downregulated), including the HSP gene *hsp-12.3* (Supplementary Fig. 4e-f).

**Figure 3:**
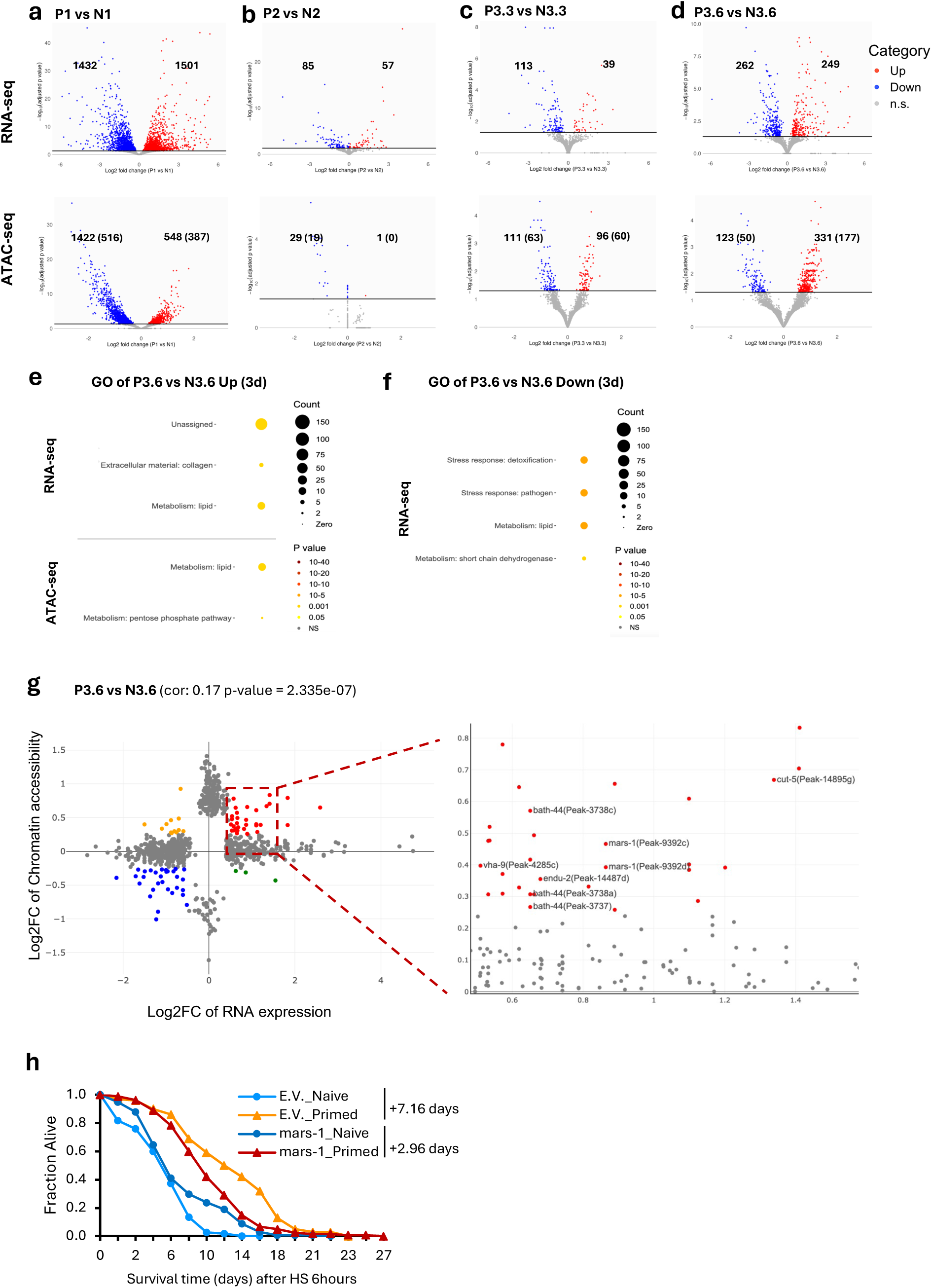
Differences in gene expression or chromatin accessibility in primed and naive worms likely contribute to the protective effects of heat hormesis. Volcano plots display differential gene expression (top panels) and chromatin accessibility (bottom panels) between primed and naïve groups at the indicated timepoints: after priming **(a),** after recovery **(b)**, after a 3-hour HS **(c),** and after a 6-hour HS **(d).** Red and blue indicate significantly up-and downregulated genes or peaks, respectively (adjusted p < 0.05; threshold shown as a black horizontal line). The numbers of significant genes or peaks (with associated genes in brackets) are indicated. Wormcat GO enrichment analysis for genes upregulated **(e)** or downregulated **(f)** in primed compared to naive groups after a 6-hour HS. Details are provided in the *source.data* and Supplementary Data 3. **(g)** Scatter plot displays genes with significant changes identified in RNA expression or chromatin accessibility after a 6-hour HS, with correlated changes between RNA-seq and ATAC-seq data highlighted according to defined criteria: log2FC RNA expression > 0.5, <-0.5; log2FC Chromatin accessibility > 0.25, <-0.25. The area showing genes upregulated in both RNA expression and chromatin accessibility (red) is zoomed out in the right panel. The five candidate genes (and their associated peaks) selected for RNAi testing are labeled. **(h)** Thermal-resistance was assessed based on the survival of *glp-1(ts)* worms treated with the indicated RNAi or empty vector (E.V.) after being subjected to our heat hormesis regimen and challenged with a 6-hour HS. Survival curves represent combined data from three independent experiments (N=3). The survival extension day indicates the mean increase in survival time (in days) of the primed group relative to the naïve group. Details can be found in Supplementary Data 4.

Intriguingly, despite minimal differences observed between primed and naive worms post-recovery, significant differences in both RNA expression and chromatin accessibility were detected between the two groups after HS of 3 or 6 hours (Fig. 3c, d; *source.data*), suggesting an underlying molecular “memory” of the earlier heat priming. As expected, the RNA expression and chromatin accessibility differences between primed and naive worms after 3 or 6 hours of HS showed a good degree of overlap (Supplementary Fig. 4a). Consistent with the phenotypic observation that primed worms showed greater survival advantage after a 6-hour HS compared to a 3-hour HS (Fig. 1d-e), we detected substantially more RNA expression and chromatin accessibility differences between primed and naive worms after a 6-hour HS (Fig. 3c-d). Specifically, 249 and 262 genes showed significantly up-or down-regulated RNA expression in the primed compared to the naive group (Fig. 3d). Additionally, 331 peaks (associated with 177 genes) and 123 peaks (associated with 50 genes) were significantly up-or down-regulated in the primed group (Fig. 3d). Among the genes with significantly upregulated expression (249) or more open chromatin (177) in the primed group, lipid metabolism and collagen were the significantly enriched GO term (Fig. 3e); Among the genes with significantly downregulated expression (262), detoxification stress response, pathogen stress response, and lipid metabolism were among the significantly enriched GO terms (Fig. 3f) (Supplementary Data 3). We conducted additional clustering analysis to assess the temporal trajectories of RNA expression across our hormesis regimen between the primed and naive groups and came to a similar conclusion that most priming-induced gene expression changes are transient, but also highlighted different RNA expression dynamics between the primed and naive groups (Supplementary text; Supplementary Fig. 4c-d).

### WT exhibits patterns of RNA expression and chromatin accessibility dynamics across heat hormesis similar to those of germlineless mutant

To investigate whether our findings are broadly relevant, and not unique to the *glp-1(ts)* mutant, we performed a parallel multiomic analysis using WT N2 strain. WT worms showed transcriptomic and chromatin accessibility dynamics highly consistent with those observed in *glp-1(ts)* mutants across the heat hormesis regimen. In WT, we again observed that the priming-responsive changes were largely restored following recovery (Supplementary Fig. 5a-b, 6b), and that different levels of heat stress induced largely distinct transcriptomic and chromatin accessibility profiles (Supplementary Fig. 5c-d). Together, these results indicate that WT and *glp-1(ts)* share similar multiomic responses, supporting that some of the molecular changes underlying heat hormesis are broadly generalizable. Interestingly, we detected fewer differentially expressed genes in WT overall, consistent with the possibility that dynamic changes in the reproductive germline could obscure somatic responses to heat hormesis. Thus, while the WT data confirm the generality of our findings, the *glp-1(ts)* background provides a clearer readout of somatic regulation that confers thermal-resistance and therefore serves as the primary dataset for identifying candidate regulators of heat hormesis.

### Multiomic data unveil putative new regulators of thermal-resistance induced by heat hormesis

To assess whether the differential molecular changes between primed and naive worms contributed to the protective effects induced by mild heat stress priming, we used two strategies to select candidate genes and tested their functional relevance in thermal-resistance using the regimen described above (Fig. 1d).

The first strategy prioritized genes that showed overlapping upregulation in both RNA expression and chromatin accessibility between primed and naïve worms after HS for 6 hours (P3.6 vs N3.6) (Fig. 3g; *source.data*). Although some of the strongest overlapping hits (top right-most dots in Fig. 3g) could not be tested due to a lack of available RNAi constructs in our library, we tested five candidates (*mars-1*, *endu-2*, *bath-44*, *vha-9*, *cut-5)* guided by their annotated gene functions. Among these, only *mars-1* knockdown significantly compromised the survival advantage of primed worms. *mars-1*, the ortholog of human MARS1, encodes a cytoplasmic methionyl-tRNA synthetase (MetRS) that catalyzes methionine attachment to its cognate tRNA, an essential step in protein synthesis. Specifically, priming induced a reduced survival benefit in worms treated with *mars-1* RNAi (mean survival extension of 2.96 days) compared to those treated with empty vector control (E.V.) (mean survival extension of 7.16 days, Fig. 3h; Supplementary Data 4). Interestingly, in naive worms, *mars-1* knockdown results in increased survival after HS compared to control (Fig. 3h). This is consistent with previous reports showing that *mars-1* RNAi enhances oxidative stress resistance and extends lifespan in *C. elegans*(31).

The second strategy focused on predicting the transcription factors that regulate the observed molecular changes, with the goal of identifying regulators that play a role in heat hormesis. We conducted motif analysis using significantly changed peak regions identified from ATAC-seq and the promoter regions of significantly changed genes from RNA-seq. The analysis revealed three groups of significantly enriched motifs: i) Motifs that were enriched based on both ATAC-seq and RNA-seq results; ii) Motifs enriched based on only RNA-seq or (iii) only ATAC-seq results (Table 1; *source.data*). From the list of enriched motifs (Q < 0.05) (source.data), we selected candidate factors that (1) were uncovered across the different datasets, (2) had available RNAi constructs, and (3) had prior evidence linking them to stress response or longevity, for further functional testing.

**Table 1:** Candidate transcription factors predicted by motif analysis and tested by RNAi The table shows the transcription factor candidates predicted by motif analysis that have been tested by RNAi, with at least two independent experiments. Promoter regions of genes associated with significant changes in RNA expression from RNA-seq and DNA regions spanning significantly changed peaks from ATAC-seq were subjected to motif analysis. Motif candidates were primarily selected from the JASPAR database, except for the HSF-1 motif, which is from CIS-BP database. Motifs identified from RNA-seq data were primarily enriched in the 500bp upstream promoter region of the TSS, unless indicated as 2000bp. The complete list of all significantly enriched motifs (Q value < 0.05) is shown in *source.data*. The table is color-coded into three groups: Motifs enriched based on both ATAC-seq and RNA-seq results (pink); Motifs enriched based on only RNA-seq (green); Motifs enriched based on only ATAC-seq results (blue). In addition to HSF-1, three candidates from each group were selected for RNAi screening if they were repeatedly identified in different significant lists, RNAi constructs were available, and the candidate factor had been implicated in stress response and longevity according to the literature.

| Candidates | P1 vs NC | P3.6 vs P2 | (P1 vs NC) & (P3.6 vs P2) intersect | P1 vs N1 | P3.6 vs N3.6 |
| --- | --- | --- | --- | --- | --- |
| cebp-1 | RNA: up | RNA: up (2000bp) |  | RNA: up |  |
|  |  | ATAC: up | ATAC: up | ATAC: up |  |
| snpc-4 |  | RNA: up |  |  |  |
|  | ATAC: up |  |  | ATAC: up |  |
| hlh-30 | RNA: down (2000bp) | RNA: up (2000bp) |  |  |  |
|  | ATAC: down | ATAC: up |  |  | ATAC: up |
| hsf-1 (CIS-BP) |  | RNA: up | RNA: up |  |  |
|  |  | ATAC: up, down |  |  |  |
| elt-2 | RNA: down |  |  | RNA: down | RNA: down |
| elt-6 | RNA: down |  |  | RNA: down | RNA: down |
| pqm-1 | RNA: down |  |  | RNA: down | RNA: down |
| dpy-27 | ATAC: down |  |  | ATAC: down | ATAC: down |
| fos-1 |  | ATAC: up |  |  |  |
| atf-7 |  | ATAC: up |  |  |  |

Among the significantly enriched motifs identified based on genes showing significantly differential RNA expression and chromatin accessibility after HS, HSF-1, SNPC-4, CEBP-1, and HLH-30 stood out (Table 1, pink section). Specifically, the motif of HSF-1, a master regulator of the heat shock response, was enriched among the genes that showed upregulated RNA expression and chromatin accessibility after priming and HS, as well as those with downregulated chromatin accessibility post HS, perhaps reflecting the transient nature of some of the HSF-1 regulated gene expression(32). SNPC-4, a component of the small nuclear RNA-activating complex, showed motif enrichment among genes upregulated after HS and regions with greater accessibility after priming. The motif of CEBP-1, a conserved bZIP transcription factor, was enriched among genes upregulated after priming. In contrast, the motif of HLH-30, the *C. elegans* ortholog of TFEB, was enriched among those downregulated after priming. RNAi knockdown confirmed that *hsf-1* is essential for heat hormesis in *glp-1(ts)*, as it completely abolished the enhanced survival to HS of the primed worms (Fig. 4a). Intriguingly, knocking down *snpc-4* also significantly compromised the survival advantage induced by priming in *glp-1(ts)* (Fig. 4b). However, no effects were detected with knockdowns of *cebp-1* or *hlh-30* under our hormesis regimen (Supplementary Data 4).

**Figure 4:**
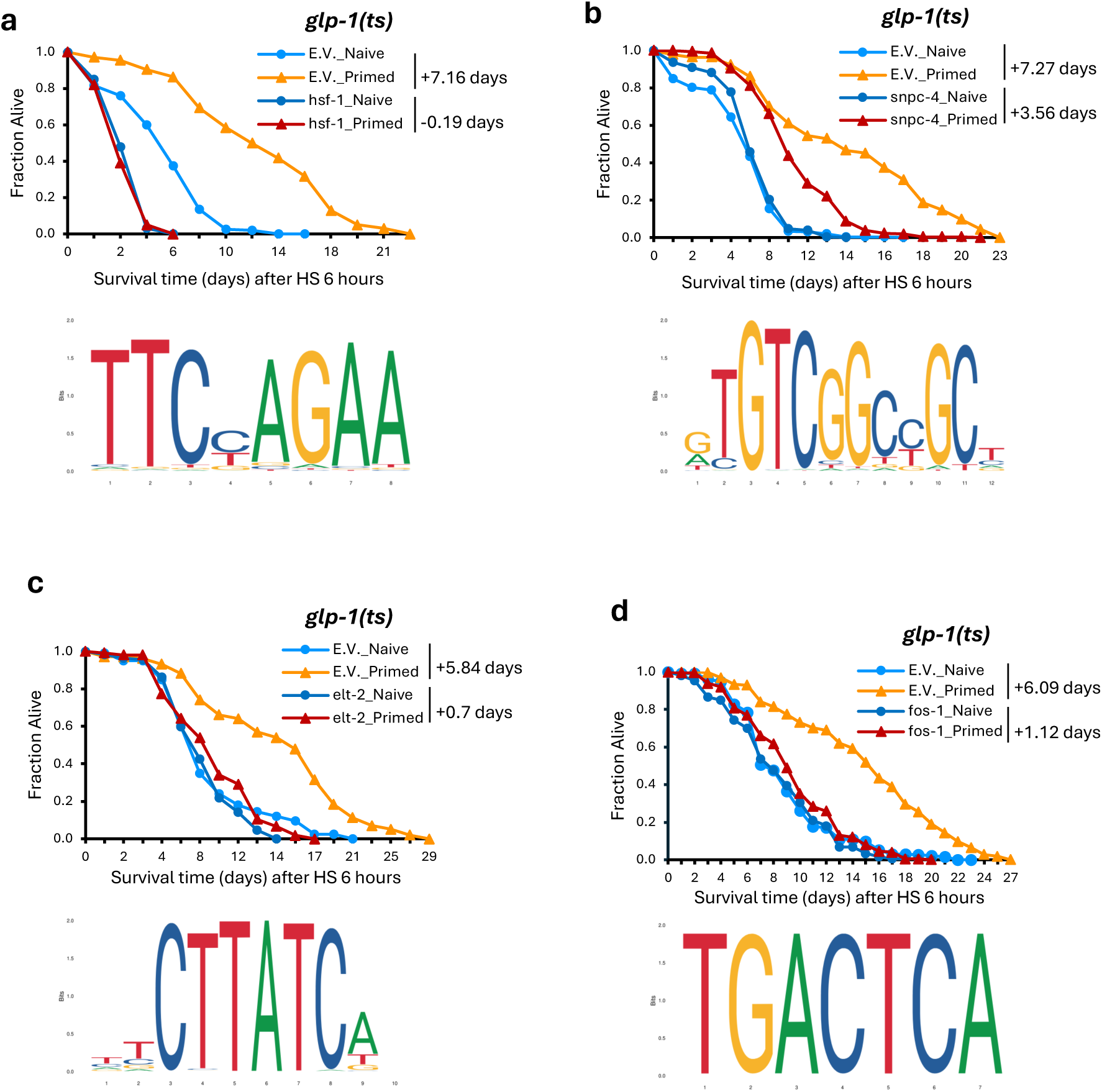
Motif analysis unveils regulators of heat hormesis. Thermal-resistance was assessed based on survival of *glp-1(ts)* mutant worms treated with the indicated RNAi and after being subjected to our hormesis regimen and challenged with a 6-hour HS. Survival curves represent combined data from multiple independent experiments (N) for *glp-1*_Naive or *glp-1*_Primed treated with empty vector (E.V.) control RNAi or *hsf-1* RNAi **(a; N=3)**, *snpc-4* RNAi **(b; N=4)**, *elt-2* RNAi **(c; N=2),** *fos-1* RNAi **(d; N=4)**. RNAi targeting *hsf-1***(a)** and *elt-2* **(c)** completely abolished the enhanced survival to heat in primed groups. For the *snpc-4* (b) and *fos-1* **(d)** RNAi, which partially compromised the enhanced survival to heat in primed groups, the survival extension rate is calculated, and the increase in mean survival days in primed worms relative to naive worms is shown. Details in Supplementary Data 4.

The motifs revealed by differential RNA-seq data included GATA transcription factors, ELT-2 and ELT-6, and a zinc finger transcription factor, PQM-1 (Table 1, green section; *source.data*). All three motifs were enriched among genes that showed downregulation after both priming and HS in the primed compared to naive groups. Interestingly, RNAi knockdown of *elt-2* completely abolished the heat shock survival advantage in primed *glp-1(ts)* worms (Fig. 4c), while knockdowns of *elt-6* and *pqm-1* had no effect (Supplementary Data 4).

Several motifs were revealed based on differential ATAC-seq data (Table 1, blue section; *source.data*). Specifically, DPY-27, a homolog of the condensin-like protein and subunit of the dosage compensation complex, had its motif enriched in peaks downregulated after both priming and HS in the primed compared to the naive groups. Additionally, FOS-1, a c-Fos ortholog and AP-1 component, and ATF-7, a bZIP transcription factor, both had motifs enriched among peaks upregulated after HS. We observed that RNAi knockdown of *fos-1* consistently compromised the heat survival advantage of primed *glp-1(ts)*, but not the other candidate factors (Fig. 4d; Supplementary Data 4). Together, these findings identify HSF-1, SNPC-4, ELT-2, and FOS-1 as putative mediators of the thermal-resistance phenotype induced by our heat hormesis regimen in *glp-1(ts)*.

We next tested whether the new regulators of heat hormesis uncovered using *glp-1(ts)* also play similar roles in WT. Interestingly, RNAi knockdown of *hsf-1, fos-1*, *snpc-4*, and *mars-1* compromised the thermal resistance benefits of primed worms in the WT background (Supplementary Fig. 7a, c, d, f; Supplementary Data 5), suggesting that these regulators are broadly important for mediating the beneficial effects of heat hormesis. We note that *hsf-1* is critical for survival post HS in both naive and primed WT worms, as *hsf-1* RNAi knockdown drastically shortened their survival post HS. However, *hsf-1* RNAi knockdown did not completely eliminate the priming-induced survival benefits in WT (Supplementary Fig. 7a), unlike in *glp-1(ts)* (Fig. 4a). Similarly, RNAi knockdown of *elt-2* did not impair priming-induced thermal-resistance in WT (Supplementary Fig. 7b), unlike in *glp-1(ts)* (Fig. 4c). In contrast, although *dpy-27* RNAi showed variable effects in the *glp-1(ts)* mutant, its knockdown consistently attenuated priming-induced thermal-resistance in WT (Supplementary Fig. 7f; Supplementary Data 5). Together, these results highlight several putative regulators of heat hormesis with major effects in both WT and *glp-1(ts)*, while also point to factors with roles in specific genetic backgrounds.

### Transient germline defects and preserved HSP inducibility may underlie lifespan extension induced by heat hormesis

Heat hormesis has been widely recognized for its longevity benefits, and previous studies have reported that WT worms exposed to the priming regimen used here (30°C for 6 hours on day 1 of adulthood) exhibited a longer lifespan(6,16). We confirmed this effect in WT; however, interestingly, the same regimen did not further extend lifespan in the long-lived *glp-1 (ts)* mutant (Fig. 5a; Supplementary Fig. 8a).

**Figure 5:**
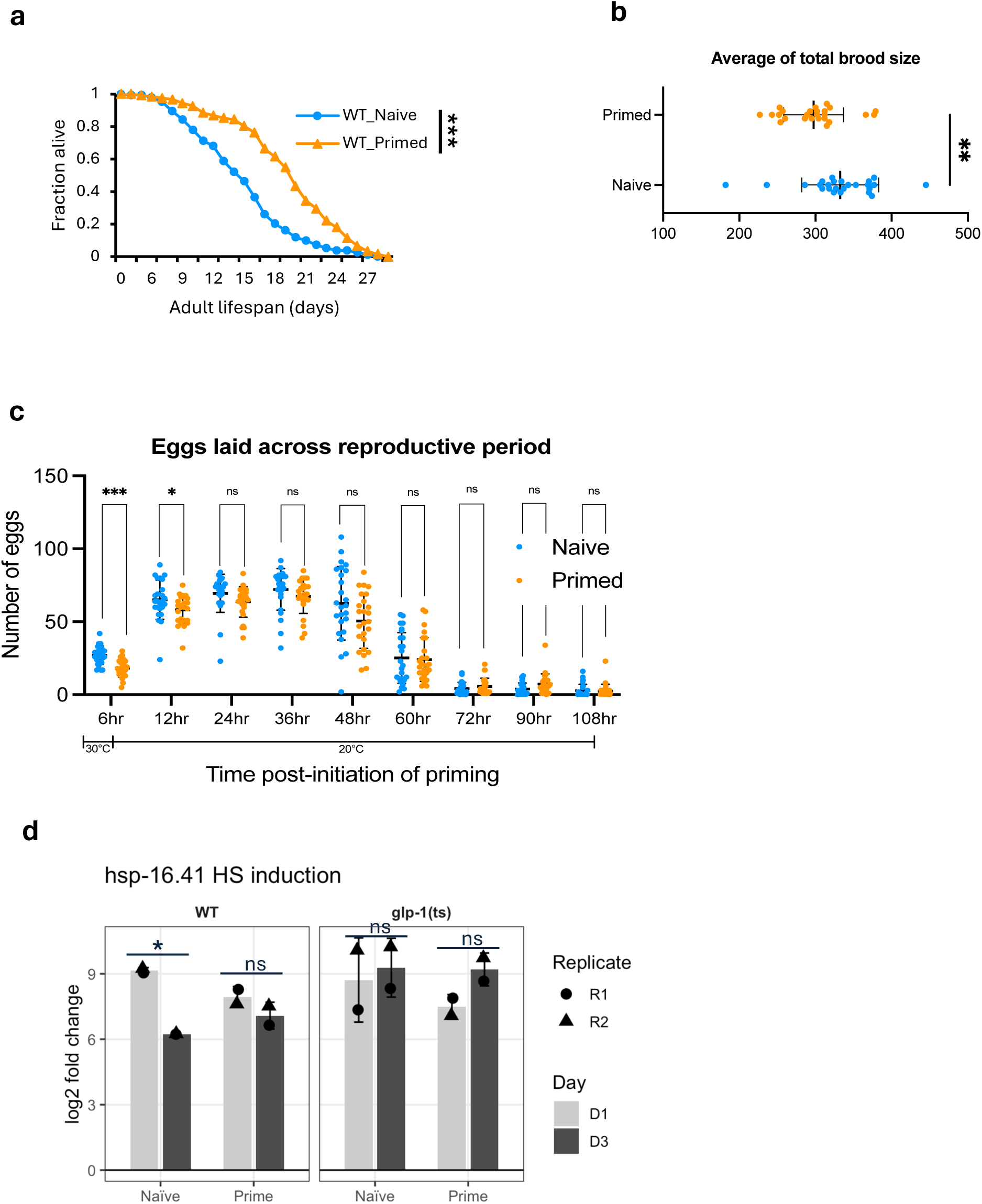
Heat priming induced lifepan extension, transient germline defects, and preserved HSP inducibility in WT. **(a)** Lifespan of WT worms with or without priming was assessed at 20 °C. Survival curves for WT_Naive and WT_Primed are shown. Data represent combined results from two independent experiments (N = 2). WT worms were maintained at 20 °C throughout their life cycle. **(b)** Average of total brood size, calculated from egg counts at nine time points across the reproductive period of individual worms, was shown in naive and primed groups. **(c)** Brood size was counted starting at the onset of priming (30°C for the primed group, 20°C for the naive group), with subsequent counts every 12 hours at 20°C. **(b-c)** were performed in WT with N=3 biological replicates and a total of 25 worms. **(d)** RNA induction of hsp genes (*hsp-16.41*) was quantified as log(2) fold change (HS/before HS) using normalized RNA-seq counts from two biological replicates (R1, R2). Log-rank test was used to compare mean lifespan in **(a).** Two-tailed unequal variances t-tests were performed in **(b–c)**, and paired t-tests were used in **(d).** * p < 0.05; ** p < 0.01; *** p < 0.001. Details are provided in the *source.data*.

To investigate this difference, we first examined reproductive function, the major distinction between the two genetic backgrounds. Priming resulted in a small but significant reduction in total brood size in WT (Fig. 5b), which was due to a significant decrease in laid embryos during the priming period (and the embryos laid during that period were also inviable), and a residual defect within 12 hours after priming, with egg laying and embryo viability returned to normal levels thereafter (Fig. 5c; Supplementary Fig. 8b).

To further understand how heat priming might impact WT worms, we compared the RNA expression changes induced by priming in WT with publicly available RNA expression profiles using the WormExp tool (33). Interestingly, for the priming-induced upregulated genes, germline stem cell removal-associated genes and *glp-1(ts)-*upregulated genes ranked among the most significantly enriched categories (*source.data: WormExp*). This finding corroborates the broodsize results and indicates that germline proliferation is transiently suspended during heat priming.

Previous research showed that WT worms exhibit an attenuated heat stress response during early adulthood, evidenced by substantially lower induction of hsp genes, including *hsp-16* and *hsp-70*, in day 2 compared to day 1 adults in response to HS (34). This is proposed to reflect maximizing germline activity rather than somatic stress responses during reproduction. To test whether heat priming affects this regulatory step, we examined the inducibility of hsp genes during early adulthood in primed and naive WT and *glp-1(ts)* worms using RNAseq data of day 1 or day 3 adult worms after HS. Our data confirmed reduced hsp induction in WT at day 3 relative to day 1, consistent with previously published data (34), while *glp-1(ts)* did not show this early-adulthood decline (Fig. 5c). Strikingly, we found that priming reversed this decline in hsp gene induction in WT, with primed day 3 adults maintaining hsp induction at levels comparable to day 1 adults (Fig. 5c; source.data). These results suggest that heat priming preserves the inducibility of the heat shock response during the reproductive period in WT.

## Discussion

Our study provides a comprehensive view of the transcriptomic and chromatin accessibility dynamics across a heat hormesis regimen in both WT and germlineless *glp-1(ts)* mutant animals, revealing molecular evidence of dose-dependent effects in which varying levels of heat stress elicit distinct molecular responses. Although priming-induced RNA expression and chromatin accessibility changes largely restore after a recovery period, the earlier stress exposure leaves distinct molecular imprints with lasting physiological consequences. Notably, our comparative analysis of WT and *glp-1(ts)* animals demonstrates that germline responses during heat priming profoundly impact long-term physiology, resulting in significant extension of mean and median lifespan. Importantly, we identify several highly-conserved regulators of heat hormesis: some act broadly across both genetic backgrounds, whereas others function in a context-dependent manner. These findings provide a molecular framework linking transient stress exposure and long-term physiological benefits, with important implications for understanding how mid-life stress management can promote healthy aging.

### Molecular evidence of dose-dependent effects

Our combined RNA-seq and ATAC-seq strategies proved complementary in uncovering molecular changes across heat hormesis. At both the level of individual genes showing changes in RNA expression and chromatin accessibility and at the level of biological functions enriched among various gene sets, our data provide compelling evidence that different levels of heat stress induce differential molecular responses. This aligns with the foundational concept of hormesis, in which a stressor generates dose-dependent effects (3,35). Since we conducted a time-series analysis, it is possible that the dose-response changes we observed are influenced by the developmental stage of the worms subjected to priming versus those subjected to heat shock. However, comparisons with published studies in which L4 or day 1 adult worms are exposed to 35°C heat shock (29,30) reveal substantially greater overlap with our heat shock response (in day 2 worms) than with the priming response (in day 1 worms) (Supplementary Fig. 3d; Supplementary Fig. 5g). This indicates that developmental timing plays a relatively minor role in the stress-induced molecular responses in our study.

### Modest correlation between RNA-seq and ATAC-seq

We observed a modest correlation between the RNA-seq and ATAC-seq datasets (Supplementary Fig. 2c), which is consistent with previous findings that ATAC-seq and RNA-seq results often do not correlate (36), reflecting different modes of gene regulation that may not be correlative. First, changes in chromatin accessibility and gene expression may occur at different times following heat stress, leading to temporal discordance between the two datasets. In particular, RNA-seq measures steady-state mRNA levels, which do not necessarily reflect active transcription at the time of sampling. In contrast, chromatin accessibility changes may more closely correlate with transcriptional activity. Second, not all gene expression changes are directly driven by alterations in chromatin accessibility. For example, transcription factors binding and local changes in chromatin could have consequential effects on gene transcription, without detectable changes in chromatin accessibility as accessed by ATAC-seq. Third, additional layers of regulation, such as post-transcriptional RNA modifications, RNA decay, or stress granule-mediated mechanisms, may also contribute. Finally, bulk sequencing approaches likely obscure tissue-or cell-type–specific responses, further contributing to the limited overall correlation between RNA-seq and ATAC-seq profiles. Future follow-up analyses, guided by limitations suggested above, including single-cell transcriptomics and genomic strategies that can distinguish immediate transcriptional responses to post-transcriptional regulation, will likely provide a more in-depth understanding of the gene regulatory landscapes across heat hormesis.

### New regulators of heat hormesis

Our multiomic data, coupled with functional screens, provided critical entry points for identifying candidate regulators of heat priming-induced thermal-resistance (Fig. 6). **HSF-1** is the master transcription factor that is essential for mounting a heat shock response (37,38) and its emergence from our analyses is a strong proof-of-principle that our investigative strategy is effective in uncovering bona fide regulators of the heat stress response. An intriguing new finding from our data is that HSF-1 appears to act somewhat differently in mediating heat priming-induced thermal-resistance in WT vs. germlineless *glp-1(ts)* strains, where it is completely required in *glp-1(ts)* but only partially required in WT. Going forward, it will be particularly important to understand the spatial regulation of HSF-1, which may illuminate how its role in mounting a heat shock response in different cells can impact overall physiological outcomes of heat hormesis. Similarly, **ELT-2**, a GATA transcription factor, is critical for intestinal development and immunity in *C. elegans* (39–41). Prior work demonstrated that ELT-2 is required for enhanced thermal tolerance in HSF-1-deficient worms (42), suggesting the possibility that ELT-2 and HSF-1 act in complementary pathways. We found that ELT-2 is essential for heat priming-induced thermal-resistance in *glp-1(ts)* mutants but dispensable in WT, highlighting a possible cross-tissue interaction between the germline and the intestine and positioning ELT-2 as a context-dependent regulator of hormesis. In contrast, we found that **DPY-27**, a homolog of human SMC4, a condensin-like subunit of the dosage compensation complex (DCC), plays a consistent role in mediating thermal-reistance of primed WT but not *glp-1(ts)*. In *C. elegans*, the DCC contributes to the formation of topologically associated domain (TAD) boundaries that regulate chromosome-wide gene expression, and elimination of DCC-dependent TADs on the X chromosome has been shown to reduce heat tolerance and shorten lifespan (43,44).

**Figure 6:**
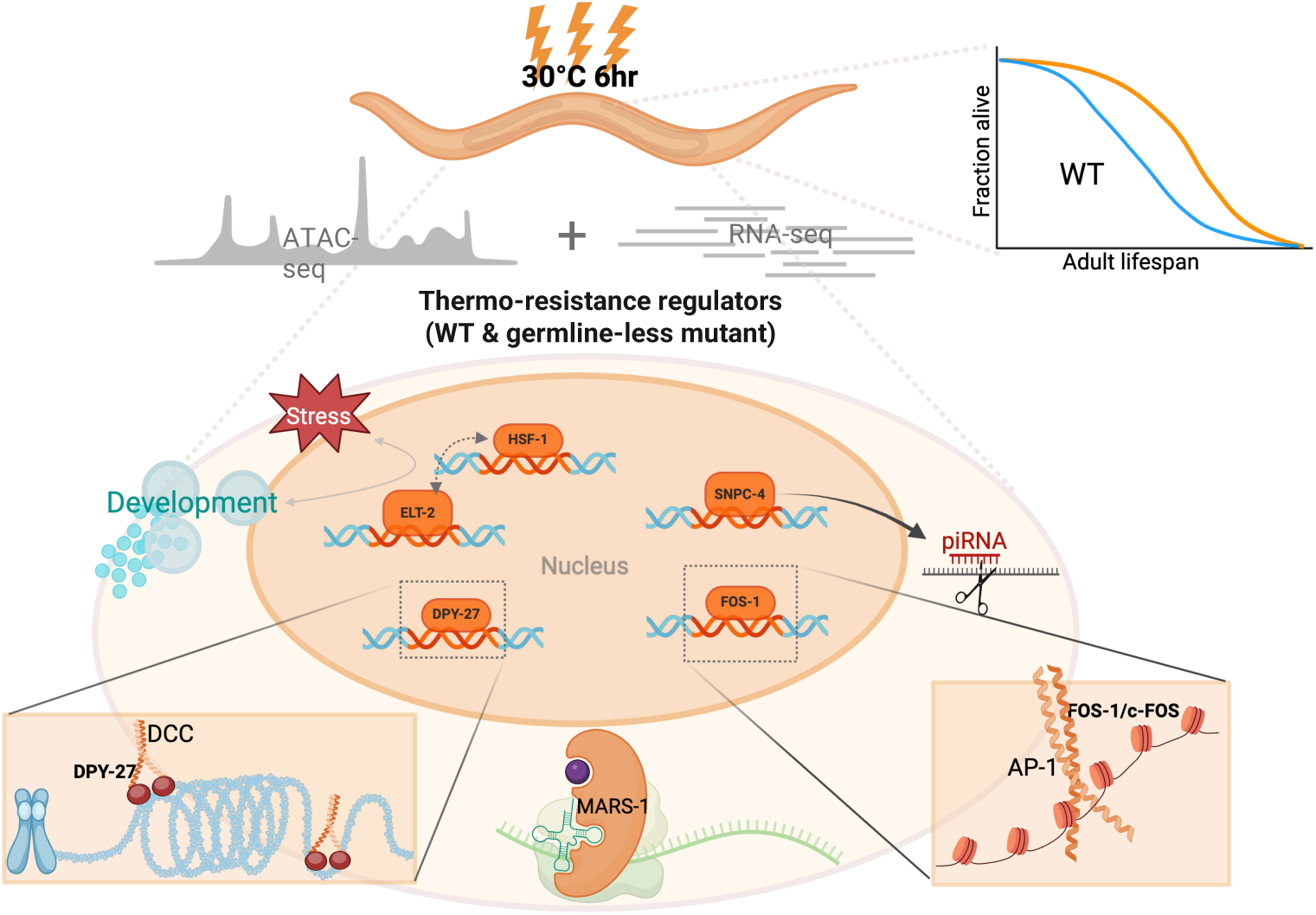
Heat hormesis engages specific regulators to induce thermal-resistance in WT and *glp-1(ts)* mutant. Our study reveals that heat hormesis promotes longevity in WT and induces thermal-resistance in both WT and *glp-1(ts)* mutant animals. Our multiomic data lead to the identification of several key regulators of heat hormesis, all of which are evolutionarily highly conserved, and participate in different regulatory steps of gene expression. HSF-1 is the master transcription factor of heat shock response and its emergence from our analyses proves our investigative strategy is effective. FOS-1 points to a potential role for the AP-1 pioneer transcription factor complex in encoding heat hormesis memory through chromatin remodeling. ELT-2 suggests an interaction between the germline and the intestine in stress adaptation. DPY-27 suggests a connection between the dosage compensation complex (DCC)-mediated chromosome architecture and heat stress management. HSF-1, ELT-2, and DPY-27 regulate heat hormesis differently in worms with or without germline. SNPC-4 implicates a role of piRNA-mediated post-transcriptional regulation in stress responses. MARS-1 implicates a role of methionine incorporation during protein synthesis in heat hormesis. The graphic was created with BioRender.com.

**FOS-1**, the *C. elegans* ortholog of c-Fos, a subunit of the activator protein-1 (AP-1) complex, is critical in mediating heat priming-induced thermal-resistance in both WT and *glp-1(ts)*. Mammalian AP-1 is well-characterized to regulate diverse processes, including stress responses (45), and *C. elegans* FOS-1 is known to regulate hyperosmotic stress (46,47). Importantly, with aging, mammalian AP-1 progressively shifts its occupancy from chromatin sites linked to developmental genes to those involved in stimuli and stress responses, resulting in remodeling of chromatin accessibility due to its pioneer transcription factor activity (48,49). The age-dependent AP-1 redistribution on the chromatin was speculated to reflect a type of “epigenetic memory”, which may align with our findings, where FOS-1 DNA motif is specifically enriched in more open chromatin regions in primed worms, suggesting its role in maintaining stress memory through chromatin opening in heat hormesis. This finding corroborates recent studies highlighting the roles of the SWI/SNF chromatin remodeling complex and the histone acetyltransferase CBP-1 in maintaining persistent induction of innate immune genes, thereby mediating stress adaptation and longevity following early-life heat exposure(21).

**SNPC-4,** a subunit of the small nuclear RNA-activating complex (SNAPc), which is essential for piRNA transcription in *C. elegans* (50,51). Interestingly, previous studies suggest that high temperatures suppress piRNA biogenesis (52). Moreover, consistent with a possible role of piRNAs in heat hormesis, *prg-1*, which encodes a Piwi-family Argonaute protein that binds piRNAs (53), is among one of the few genes that showed persistent upregulation through heat hormesis (*source.data: Fig. 3a-d_RNA-seq_glp-1(ts)* – highlighted in yellow). These findings implicate post-transcriptional regulation in heat hormesis. **MARS-1** is a conserved methionine tRNA synthetase. *mars-1* knockdown enhances thermal-resistance of naïve worms, and heat priming only confers marginal further protection. Interestingly, human MARS1 mutations have been implicated in chronic activation of the integrated stress response (54), suggesting conserved roles in stress response from worms to humans. The identification of MARS-1 from our study implicates a possible link between methionine incorporation during protein synthesis and priming-induced protective effects.

Together, the putative new regulators of heat hormesis identified here point to different steps of gene expression regulation in mediating the beneficial effects of heat hormesis. These regulators are highly conserved, and their further characterization promises to unveil mechanistic principles that can be broadly relevant to stress adaptation and healthspan regulation.

### Germline-dependent mechanisms of heat hormesis

Our complementary studies in WT and *glp-1(ts)* led to the intriguing finding that heat hormesis likely extends lifespan through a germline-dependent mechanism. Multiple lines of evidence support this conclusion: First, heat priming only extends lifespan in WT, not in germlineless *glp-1(ts)* mutants.

Second, heat priming transiently disrupts reproduction. Third, heat priming induces gene expression changes in WT animals that strongly resemble those associated with germline ablation. Fourth, heat priming preserves hsp gene inducibility upon HS during early adulthood, similar to the effect of *glp-1(ts)* mutation. Notably, while germline stem cell ablation or its genetic mimic, such as the *glp-1(ts)* mutation, is well-established to extend lifespan in *C. elegans* (55) and other diverse species (56–58), our findings demonstrate that temporary, transient disruption of germline activities in response to heat stress produces a prolonged and significant effect on physiological effects. Furthermore, our data reveal that specific regulators of heat hormesis, such as HSF-1, ELT-2, and DPY-27, regulate stress adaptation in a genetic background-dependent manner. These results have broader implications: whereas hormetic stress can enhance stress resilience and longevity, its benefits may be intertwined with trade-offs in reproductive or developmental functions. Future research to further investigate these complex relationships at the molecular level, especially with spatial resolution, will be critical for developing stress management strategies that promote healthy aging without compromising quality of life in the short term.

We additionally observed that our heat hormesis regimen primarily extends mean and median lifespan rather than maximum lifespan and induces a slight “squaring” of the survival curve (Fig. 5a; Supplementary Fig. 8a), indicating a significant reduction of early-and mid-life mortality. Importantly, this pattern resembles findings from worms experiencing a higher oxidant state during development (59), suggesting that lowering early-and mid-life mortality and improving mean and median lifespan, likely reflecting improved healthspan, may represent a universal signature of stress hormesis, independent of the specific stressor. Elucidating how different stressors confer common as well as unique stress responses and adaptations will be a critical area of future research that will inform potential health-promoting interventions via stress exposure and management.

## Methods

### *C. elegans* maintenance

Unless otherwise specified, worms were maintained on 6-cm NGM plates seeded with 0.2 ml of a five-times concentrated overnight culture of streptomycin-resistant OP50 bacteria (live OP50). Killed OP50 bacteria were prepared by resuspending live OP50 in LB containing 100µg/ml Carbenicillin and 15µg/ml Tetracycline twice, then concentrating it twentyfold. The bacteria incubated with the antibiotic on a rocker for at least 30 minutes before use. Killed OP50 was used to mitigate bacterial infections in *glp-1 (ts)* mutant strains (*e2144* temperature sensitive allele) during survival assays. The N2 strain was typically kept at 20°C, while the *glp-1 (ts)* mutant strain was maintained at 16°C unless noted differently.

### Growth of synchronized worms on a small scale for phenotypic experiments

#### For the wildtype N2 strain

Several plates were set up with 4-5 gravid adults per plate for egg laying at 20°C for 3-4 hours. After egg laying, the gravid adults were removed, and the eggs were incubated at 20°C for approximately 65 hours. The synchronized gravid adults were then distributed to fresh plates, approximately 30 per plate, before the experiments began. Priming was performed at this time when the N2 worms had just reached the gravid adults of their Day 1 (D1) adult stage. **For the *glp-1(ts)* mutant strain:** Several plates were set up with 6-8 gravid adults per plate for egg laying at 16°C for 4-5 hours. The gravid adults were removed after egg laying, and the eggs were incubated at 25°C for approximately 48 hours until they became young adults and were transferred to 20°C (“Young adult” refers to the developmental stage immediately after the L4 stage, in which the worm has molted into a reproductive adult but has not yet begun laying a large number of eggs). After incubating overnight at 20°C (approximately 15-17 hours), the synchronized adults were distributed to fresh plates with Killed OP50, approximately 30 per plate, before starting experiments. Priming was performed at this time when sterile *glp-1(ts)* worms were approximately at their D2 adult stage. The wildtype N2 was sometimes conducted in parallel as a positive control, following the same maintenance protocol as *glp-1(ts)*.

### Priming and Heat shock (HS) experimental setup

Worms were subjected to incubators with setting pre-adjusted to target temperatures (30°C for 6 hours for priming, and specific temperatures and duration for HS depending on the experiment). During incubation, the lids of the plates were replaced with those having manually drilled holes, and the plates were wrapped with parafilm on the sides. The plates were placed on a metal rack in the incubator with agar side facing down. This setup aimed to allow worms to reach the target temperature more efficiently and to maintain consistent temperature and humidity inside the plates. After temperature treatments, the parafilm was removed, and the lids were switched back to the normal ones without holes before being moved back to 20°C.

### Thermal-resistance assay

To explore how priming affected thermal-resistance, defined as regaining motility after exposure to the indicated HS temperature and duration, we conducted tests. For small-scale phenotypic experiments, 30-40 worms were placed on each 6-cm plate, with at least two plates per group for each biological replicate. For large-scale experiments, approximately 1000 worms were placed on each 10-cm plate. After being subjected to HS at 35°C for 10 hours or other specified temperatures and durations detailed in the *source.data*, worms were maintained at 20°C until scoring. Scoring occurred 15 hours post-HS, with each plate being tapped 10 times consistently immediately before scoring. For small-scale experiments, worms that moved without being touched by a picker were counted, along with the total number of worms, to calculate the percentage of moving worms. For large-scale experiments, a consistently marked 1/9 area of each 10 cm plate was used to count worms that moved without being touched and the total worms within the marked area to calculate the percentage of moving worms. Three separate areas were marked and scored for each experimental group as technical replicates. Three independent experiments were conducted as biological replicates. The average percentage of moving and standard deviation (std) were calculated. 2-tailed unequal variances t-test was performed to determine if the differences between primed and naive groups were significant.

### Survival assay after HS

Synchronized worms were subjected to priming, recovery, and HS or were maintained without priming for the naive group, according to specific temperatures and durations as detailed in *source.data*. After HS, worms were scored every two days until all had died. Worms were scored as dead when they failed to respond to a gentle prod on the head by a worm picker. Worms were maintained at 20°C after HS and throughout the experiments. Data were analyzed using OASIS2 online survival analysis tool(60). The Kaplan-Meier estimator was used to re-plot survival curves in Excel, with the ‘Fraction alive’ on the y-axis and days after HS on the x-axis. Log-rank tests were utilized to determine if there were significant differences in survival times between the two groups. All survival assay experiments were conducted independently at least twice. (See *source.data* for raw data and analysis details)

### Lifespan

Synchronized worms were subjected to priming, or were maintained without priming for the naive group, at the indicated timing mentioned above and detailed in *source.data*. Worms were scored subsequently every two days until all had died. Worms were scored as dead when they failed to respond to a gentle prod on the head by a worm picker. For the wildtype N2 strain, worms were transferred to fresh plates every 1-2 days until the end of their reproductive period. The chemical 5-fluoro-2’-deoxyuridine (FUDR) was not used. For *glp-1(ts)* mutant strain, worms for both primed and naive groups were transferred to plates with Killed OP50 before the priming experiments. Data were analyzed using OASIS2 online survival analysis tool as mentioned above. The Kaplan-Meier estimator was used to re-plot survival curves in Excel, with the ‘Fraction alive’ on the y-axis and days after HS on the x-axis. Log-rank tests were utilized to determine if there were significant differences in survival times between the two groups. All lifespan assay experiments were conducted independently at least twice. (See *source.data* for raw data and analysis details)

### Reproductive function experiments

Wildtype N2 worms were maintained, synchronized at 20°C. After approximately 65 hours of development from eggs, newly matured gravid adults were distributed, one per 3.5 cm plate. The primed group was subjected to a 6-hour priming at 30°C and returned to 20°C, while Naive group was maintained at 20°C in parallel during priming period. After priming, each worm was transferred to a new 3.5 cm plate to continue laying eggs, and the eggs left on the original plate were counted. The worms were subsequently transferred to new plates every 12 hours until they stopped laying eggs (∼5days). Once counted, the eggs were incubated at 20°C to allow for hatching. Larvae were then counted 24 hours later. In total, 9 data points for both egg and larvae counts were collected. The average total broodsize was calculated by summing egg counts from 9-timepoints across the reproductive period of 25 individual single worms. The average broodsize for each time frame (6 hours for the first timepoint and 12 hours for the subsequent eight timepoints) was calculated across these 25 worms. The average hatching rate for each timepoints was calculated by dividing the number of eggs by the number of larvae for 25 worms. Hatching rates were adjusted to 1 if the values exceeded 1 to account for potential miscounts of eggs, which might be less visible than larvae during counting. Data of 25 worms were collected across three independent experiments, with each biological replicate containing around 5∼10 worms per group. (Details in *source.data)*

### RNA interference

HT115 bacteria expressing double-stranded RNA against the gene of interest were obtained from the Ahringer(61) or Vidal(62) libraries and verified by Sanger sequencing. RNAi bacteria and the control empty vector (E.V.), L4440, were cultured overnight in LB containing 100 µg/ml carbenicillin and shaken at 37°C for 3-4 hours until an optical density of 0.6-0.8 was reached. The cultures were then induced with 1mM of IPTG and continued to incubate on a shaker at 37°C for an additional 2.5 hours. After induction, the bacteria were centrifuged at 2000 rpm for 15 minutes at room temperature and concentrated 50-fold. 200µl of concentrated bacteria were seeded on 6-cm ‘RNAi plates”, which are NGM plates modified by omitting streptomycin and adding 100 µg/ml ampicillin, 15 µg/ml tetracycline, and 1mM IPTG. RNAi was administered to worms through feeding. The timing of RNAi administration varied depending on the genes, considering any developmental phenotypes caused by knocking down the gene. Details of the RNAi administration timing for each gene are documented in the Supplementary Data 4-5. To determine whether RNAi knockdown of a specific gene compromises the priming-induced phenotype in survival extension, the survival extension rate is calculated using the formula: *(Mean survival days in primed−Mean survival days in* Naïve*)/Mean survival days in naïve * 100*. This measure quantifies the increase in mean survival days in primed group relative to naïve group in each independent experiment. To statistically evaluate whether the survival extension is significantly reduced upon gene silencing, a paired two-tailed t-test is employed.

### Collection of timed samples synchronized worms on a large scale for sequencing

For *glp-1*(ts) mutant strain, five gravid adults were grown on several 6-cm NGM plates containing 0.2ml of a five-times concentrated overnight culture of streptomycin-resistant OP50 for 6 days at 16°C until the plates were full of mixture stages of worms and embryos without running out of the OP50. 1ml M9 buffer with 0.05% TWEEN20 was added into each 6-cm plate without disturbing the bacteria lawn to avoid collecting embryos. (TWEEN20 was used to avoid worms sticking on the tips during the process) M9 buffer with 0.05%TWEEN20 containing adults and larvae was then collected into centrifuge tubes. The supernatant containing larvae was removed around 30-60 sec while most of the adults settled down to the bottom of the tube. Using this strategy to separate adults and larvae by washing worms several times until the tube mostly contained adults. Fully resuspended adults in 1 ml M9 buffer with 0.05% TWEEN20 and aliquot 10 µl out to count the number of adults. Based on the counting numbers, proportionally distributed the desired amount of the adults to new plates for egg laying.

10-cm NGM plates containing 1ml of a 25-times concentrated overnight culture of streptomycin-resistant OP50 were used for growing larger amounts of worms. 1000-2000 adults were placed on 10-cm plates and incubated at 16°C for 4-5 hours for egg laying. 5ml M9 buffer was applied to wash off adult worms from the plates without disturbing the bacteria lawn. Repeated the washing step until most of the adults were removed. 5ml m9 with 0.05%TWEEN20 was applied to wash off the bacterial lawn containing embryos using the liquid force generated by several pipetting. The liquid containing embryos were then collected into centrifuge tubes and were spun down in a centrifuge for one minute at 2000 rpm. Removed the supernatant and resuspended the pelleted embryos thoroughly in My with 0.05% TWEEN20. The number of synchronized embryos was quantified by proportionally counting and around 1200 embryos were distributed per 10-cm plates. At the assigned timepoints, synchronized worms were washed off and collected into centrifuge tubes using M9 with 0.05% TWEEN20. After the worms settled in the bottom of the tubes, removed supernatant and transferred worms to 1.7ml low-binding tubes. Worms were washed by M9 buffer several (1–5) times until the supernatant was clear indicating most of the bacteria was removed. 200 worms were then aliquoted to another low-binding tube and snap-frozen in 1ml TRIZOL after removing most of the M9 buffer in liquid nitrogen for RNA-seq. The remaining 1000 worms were snap-frozen after removing most of the M9 buffer in liquid nitrogen for ATAC-seq. After snap frozen, all samples were stored at-80°C until further processing.

Five biological replicates for RNA-seq (labeled R_r1, R_r2, R_r3, R_r4, R_r5) and four for ATAC-seq (labeled A_r1, A_r2, A_r3, A_r4) were processed in two independent sequencing batches. For RNA-seq, R_r1, R_r2, and R_r3 were in the first batch, and R_r4 and R_r5 in the second. For ATAC-seq, A_r1 and A_r2 were in the first batch, and A_r3 and A_r4 in the second. Each replicate includes samples from multiple timepoints across our regimen in both primed and naive groups (Fig. 1a). Data from all timepoints have at least three biological replicates, except for T3.6, which has two biological replicates in ATAC-seq.

### RNA extraction and RNA-seq

200 worms per sample were snap-frozen in 1 ml of TRIzol, as previously described, and homogenized by alternating between thawing, vortexing, and refreezing in liquid nitrogen four times. 200 µl of chloroform was added to each tube, samples were vortexed for 15 seconds, incubated at room temperature for 3 minutes, and then centrifuged at 12,000 g for 15 minutes at 4°C. The upper aqueous layer was transferred to a fresh tube, and 500 µl of isopropanol was added to each sample. After adding 1 µl of GlycoBlue and incubating for 15 minutes at room temperature, samples were centrifuged at 12,000 g for 10 minutes at 4°C. The supernatant was discarded, and the RNA pellet was washed with 1 ml of 80% ethanol. All remaining ethanol was removed after centrifugation at 12,000 g for 10 minutes at 4°C, and the RNA pellet was air-dried for 5-7 minutes and then dissolved in 17 µl of DEPC water. The RNA was then treated with DNase to remove residual DNA, following the instructions from the TURBO DNase kit (Invitrogen AM1907). After DNase treatment, the RNA was further purified using an RNA cleanup kit (ZYMO Research R1015). RNA concentration was measured using the Qubit RNA HS Assay kit (ThermoFisher Q32851). Five biological replicates were processed as described above.

RNA-seq libraries were prepared using the QuantSeq 3’ mRNA-Seq Library Prep Kit (Lexogen FWD 015). For replicates R_r1, R_r2, and R_r3, the preparation started with 100 ng of RNA from each sample and employing 15 PCR cycles. For replicates R_r4 and R_r5, the libraries were prepared starting with 70 ng of RNA and employing 17 PCR cycles. The libraries were quantified using a Qubit DNA HS Assay kit and their quality was assessed with a Bioanalyzer. Subsequently, the libraries were submitted for single-end 86 bp sequencing on an Illumina NextSeq 500 machine. Replicates R_r1, R_r2, and R_r3 were pooled in one sequencing lane, while R_r4 and R_r5 were pooled in another, with each pool sequenced in separate runs.

### RNA-seq data analysis

#### Upstream analysis

Adaptor sequences were trimmed, and low-quality reads filtered from raw fastq files using the trim_galore function in Cutadapt (version 4.6) with settings-q 20 --fastqc. Trimmed sequencing reads from the FASTQ files were then aligned to the reference genome (ce11/WBcel235) using STAR aligner. Initially, the reference genome was indexed with the following settings: --runThreadN 8 --runMode genomeGenerate --genomeDir [genome_directory] --genomeFastaFiles [genome fasta_file] --sjdbGTFfile [gtf_file] --sjdbOverhang [read length – 1]. Alignment was performed using STAR settings: --quantMode GeneCounts --genomeDir [genome_directory] --readFilesIn [trimmed sequencing file] -- readFilesCommand zcat --runThreadN 2 --outFileNamePrefix [output_file_prefix] -- outFilterMultimapNmax 1 --outFilterMismatchNmax 2 --outSAMtype BAM SortedByCoordinate. The resulting tab-delimited text files, named with the prefix “_ReadsPerGene.out.tab”, contained counts for reads aligned to the plus strand of RNA in column 3, which is recommended and applied for 3’ RNA-seq data. Counts from each sample were aggregated to generate a matrix that was uploaded to RStudio for downstream differential gene expression analysis (Supplementary Data 8).

Total aligned reads and the percentage of uniquely mapped reads from all sequencing files were accessed. Only uniquely mapped reads were utilized for downstream analysis (*source.data*: RNA-seq QC). The correlation of aligned reads across biological replicates was calculated using Spearman correlation and visualized with a heatmap using Morpheus (https://software.broadinstitute.org/morpheus).

(Supplementary Fig. 2a-e). The color scheme of the heatmap displays a minimum value of 0.85 and a maximum value of 1.

### Differential RNA expression analysis

A matrix (Supplementary Data 12) containing raw read counts was generated and uploaded to RStudio for DESeq2 analysis. The matrix was subsetted according to timepoints and replicates for further comparison. For comparisons involving timepoints 1, 2, and 3.3, columns including five replicates (R_r1, R_r2, R_r3, R_r4, and R_r5) for samples (NC, N1, P1, N2, P2, N3.3, and P3.3) were subsetted. For comparisons involving timepoint 3.6, columns including three replicates (R_r1, R_r2, and R_r3) for samples (NC, N1, P1, N2, P2, N3.3, P3.3, N3.6, and P3.6) were used. Prior to differential analysis, read counts were normalized using the estimateSizeFactors function. The normalized counts were then filtered to retain only genes with more than 5 normalized counts in at least the minimum number of replicates per sample. To better estimate log2 Fold Change (log2FC) for genes with low counts and high dispersion, the apeglm method was employed for shrinkage during differential analysis. Genes with adjusted p-values less than 0.05 were considered significant. MA and Volcano plots were generated using methods described in the ATAC-seq analysis section (Supplementary Data 8)

### Nuclei purification and ATAC-seq

Worms, previously snap-frozen and stored in low-binding tubes, were thawed on ice. All subsequent steps were conducted on ice or at 4°C, with all materials pre-chilled to maintain nuclei integrity. An equal volume of freshly prepared Nuclei Purification Buffer (NPB)(63) (recipe available in *source.data: ATAC protocol*) was added to the worm pellet. The worms were homogenized manually five times using Pellet Pestles coated with Fetal Bovine Serum. The homogenization frequency was optimized to minimize damaging nuclei. After homogenization, worm pellets were allowed to settle for 3-5 minutes, then centrifuged at 100-200 g for 3 minutes. The supernatant, containing the nuclei, was transferred to a fresh low-binding tube. The remaining worm pellets were resuspended in an equal volume of 2X NPB and homogenized again as described. This process was repeated for multiple rounds until no visible worm pellet remained, typically requiring 7-8 rounds. The first six rounds involved homogenizing five times, with the final rounds reduced to three times to preserve nuclei integrity. After completing the homogenization, the collected supernatants were centrifuged at 100-200 g to remove debris, transferring the clean supernatant into a new tube. A subset of nuclei was stained with DAPI and examined under a fluorescence microscope to assess quality. Nuclei counts were determined using a hemocytometer with DAPI staining. For ATAC-seq, 50,000 nuclei per sample were aliquoted into a standard tube (non-low-binding) and pelleted by centrifugation at 1,000 g for 10 minutes. The pellet location was marked, and the supernatant was carefully removed to avoid contamination with bacterial DNA.

Purified nuclei were immediately subjected to the ATAC-seq procedure. Nuclei were gently resuspended in 47.5 µl Omni-ATAC buffer (composed of 2X Illumina Tagment DNA (TD) buffer, 16.5 µl 1X PBS, 0.1% TWEEN20, and 0.01% Digitonin) and mixed with 2.5µl of Illumina Tagment DNA Enzyme (TDE1) (Cat#20034197). The mixture was incubated at 37°C on a thermomixer set to 500 rpm for 0.5-1 hour. Tagmented DNA was then purified using a MinElute PCR Purification Kit (Qiagen #28004) and eluted in 10µl. The purified DNA fragments were stored at-20°C until PCR amplification (Details in *source.data*: ATAC protocol).

PCR amplification was performed using NEBNext Ultra II Q5 Master Mix (M0544S) in a total volume of 50 µl, with 9 µl of purified tagmented DNA and 25 µM primer concentration. The thermocycling protocol included an initial step at 72°C for 5 minutes, followed by 98°C for 30 seconds, and 12 cycles of 98°C for 10 seconds, 63°C for 30 seconds, and 72°C for 1 minute. Each sample utilized a common Adapter 1 primer (Ad1) and a unique index Adapter 2 (Ad2*) primer (Details in the *source.data*: ATAC protocol).

Amplified ATAC-seq DNA libraries were purified using the MinElute PCR Purification Kit (Qiagen #28004) and eluted in 25µl of warm DEPC water. Libraries were size-selected to retain DNA fragments between 100bp-600bp using Blue Pippin service, quantified using a Qubit, and quality-checked using a Bioanalyzer. Sequencing was performed on an Illumina NextSeq 500 machine, with libraries from replicates A_r1 and A_r2 pooled at a final concentration of 3.6 ng/µl for one sequencing lane, and libraries from replicates A_r3 and A_r4 pooled at a concentration of 1.96 ng/µl for another lane. Each pool was sequenced independently.

### ATAC-seq data analysis

#### Upstream analysis: From Raw data to final BAM files

FASTQ files were downloaded using the command wget-q-c-O. The paired-end FASTQ files, denoted as.R1 and.R2, were organized for processing. Adapters were trimmed using Cutadapt (version 4.6) integrated with Trim Galore, applying the parameters --gzip-nextseq 20 --cores 8 --paired. The trimmed, paired-end sequencing files were then aligned to the reference genome Caenorhabditis elegans WBcel235 (ce11) using BWA with settings-t 8-M. Following alignment, SAM files were converted to BAM format and subsequently sorted and indexed using SAMtools. To refine the BAM files further, mitochondrial DNA sequences were excluded using samtools view. Sequences listed in the Caenorhabditis elegans ce11 blacklist (version 2) were also removed using bedtools intersect. Duplicate reads were marked using Picard Tools’ MarkDuplicates and excluded with samtools view-F 0X400. The resultant BAM files, purged of mitochondrial DNA, blacklist regions, and duplicates, were indexed for downstream analysis.

### Quality control (QC) and Transcription Start Site (TSS) enrichment analysis

Quality control metrics were assessed using several tools. Aligned and uniquely mapped reads were quantified using samtools view-c, while samtools idxstats provided percentages of mitochondrial DNA (mtDNA) reads. Duplicate read percentages were determined using Picard Tools’ MarkDuplicates function and summarized in MultiQC reports (*source.data*: ATAC-seq QC). For TSS enrichment analysis, the DeepTools suite was utilized. BAM files were first converted to BigWig format with bamCoverage, which translates raw read counts into coverage tracks. This step was performed without scaling adjustments to preserve original read densities. TSS signal coverage was then calculated for regions spanning 1000 bp upstream and downstream of each TSS using the computeMatrix command (settings: reference-point-a 1000-b 1000). Heatmaps visualizing these coverage intensities were generated with plotHeatmap. All samples demonstrated enriched reads at TSS, aligning with a distinct peak at each TSS. The reference TSS data, adopted from Chen et al.(64), was based on transcription initiation cluster (TIC) ‘mode positions’ mapped to the ce10/WS220 genome. To ensure compatibility with our analyses, which utilized the ce11 reference genome, TSS data was converted using the UCSC LiftOver Tool

### Peak calling

Following quality control of the upstream analysis, final indexed BAM files were utilized for narrow peak calling. Each sample and replicate underwent peak calling independently using MACS2 (version 2.2.7.1-r9). The settings employed were-f BAMPE --bdg --SPMR --gsize ce-q 0.05 --call-summits, with the default local correction parameters set to --slocal 1000 --llocal 10000. For visualization of read coverages, the makeTagDirectory function from the Homer software suite was used to create tag.dir. Bedgraph files were then generated using the makeUCSfile function with the parameters.tag.dir-o auto-fsize 1e10-res 1-color 106,42,73-style chipseq.

### Identifying ‘consensus peaks’ and quantification of reads in peaks

To identify’consensus peaks’ for differential chromatin accessibility analysis, a three-step process was implemented:

1. **Merging read files:** BAM files from all biological replicates within each experimental group were merged using the samtools merge command. Experimental groups included NC, N1, N2, N3.3, P1, P2, P3.3 with four biological replicates each, and N3.6, P3.6 with two biological replicates each.
2. **Calling consensus peaks:** Using MACS2, consensus narrow peaks were called from a single command line that included all merged BAM files as inputs, using settings previously described.
3. **Subdividing peaks based on summits:** The identified consensus narrow peaks were further subdivided based on their summits using the splitMACS2SubPeaks.perl script (adapted from Daugherty et al.(63). This subdivision produced a total of 30,404 ‘Consensus Peaks’ with an average length of 481 bp. Details of the genomic region for the 30,404 Consensus Peaks, labeled with the PeakID syntax’4reps.HS6hr_ConsensusPeaks_peak_’, are available in *source.data: 30,404 Consensus* Peaks.

Quantification of reads in peaks: Reads within these ‘consensus peaks’ for each individual sample were quantified using the featureCounts program, facilitating subsequent analyses of chromatin accessibility variations across samples.

### QC and consistency of biological replicates

FRiP scores, representing the fraction of mapped reads in identified region of consensus peaks, were calculated for each sample using the featureCounts program and aggregated with multiqc. Across all samples, FRiP scores ranged from 31% to 49%, indicating no significant variance among biological replicates and suggesting reliable peak calling and reproducibility across different batches (Supplementary Fig. 2k). To further assess the consistency of biological replicates, genome-wide correlations of mapped reads between biological replicates were analyzed using the bedtools multicov function with a 2 kb sliding window across the ce11genome, segmented by the bedtools makewindows function. Read counts from each BAM file in each 2 kb window were logged (log10 transformation) and uploaded into RStudio. Pearson’s correlation coefficients between each replicate were calculated using the cor function in R, compiled into a matrix, and visualized using the heatmap.2 function from the gplots package in R, categorizing samples by timepoints for clarity in correlation assessment (Supplementary Fig. 2f-j).

(Supplementary Data 6)

### Differential analysis of chromatin accessibility

A matrix containing raw read counts in consensus peaks was generated and uploaded to RStudio for DESeq2 analysis. The matrix encompassed four biological replicates (A-r1, A-r2, A-r3, A-r4) for samples: NC, N1, P1, N2, P2, N3.3, and P3.3 and two biological replicates (A-r1, A-r2) for the samples: N3.6 and P3.6. (Supplementary Data 13). This matrix was loaded into RStudio and served as input for DESeq2 for differential analysis. Prior to analysis, read counts were normalized using the estimateSizeFactors function. Only peaks with more than five normalized counts in at least two samples, the minimum number of replicates in some experimental groups, were retained for analysis. To refine the estimation of log2 Fold Change (log2FC) for peaks with low counts and high dispersion, the apeglm method was utilized for shrinkage during the differential analysis. Peaks with adjusted p-values below 0.05 were determined significant. MA and Volcano plots were generated via ggplot2, where peaks significantly upregulated are marked in red and those downregulated in blue, unless labeled specifically. (Supplementary Data 8).

### Peak annotation

For annotating consensus peaks, a dataset of 42,245 accessible elements, termed Reference Elements, derived from the study by Jänes et al.(65) (elife-37344-fig1-data1-v2) was utilized. These Reference Elements, adapted using ce11 genome information (Supplementary Data 14), facilitated the annotation of consensus peaks, which involved matching consensus peaks to Reference Elements with at least a 50% overlap. This annotation process was conducted using the ‘findOverlapsOfPeaks’ function within the ‘ChIPpeakAnno’ R package (Supplementary Data 9). A total of 30,404 consensus peaks were annotated, linking them to 19,352 functional information records that included associated genes and functional elements such as promoters and enhancers (*source.data: annotated.ConsensusPeaks*).

### Motif analysis

Motif enrichment analysis was performed using the Simple Enrichment Analysis (SEA) tool from THE MEME Suite (version 5.5.7)(66). To identify motifs enriched in the promoter regions of significant genes from RNA-seq data, promoter sequences were obtained from the regions either 500bp or 2000bp upstream of the TSS following ‘Supplementary Data 10’. The input sequences comprised promoter sequences from significant genes, while control sequences were promoter sequences from non-significant genes. For motif enriched in significant peak regions from ATAC-seq data, BED files composed of significant peak regions were used as input sequences, while non-significant peak regions were used as control sequences.

The ‘JASPARA (non-redundant)-nematode2022(67)’ and ‘CIS-BP 2.0 Single Species-Caenorhabditis_elegans(68)’ motif databases were applied to identify motif enrichments. Enrichment was considered significant if the Q-value was less than 0.05. Significantly enriched motifs were selected as candidates for RNAi screening if their associated genes showed detectable expression levels in our RNA-seq data. We primarily selected motif candidates identified in the JASPARA database for our RNAi screening, as the database has undergone more functional validation compared to the motifs in CIS-BP. Similar motifs were detected in both databases, while the hsf-1 motif was found exclusively in CIS-BP and not in JASPAR. (Motif list can be found in *source.data*)

### Temporal dynamic analysis and trajectory plotting

For the temporal dynamic analysis, data input of genes with their Log2FC between prime and naive groups across timepoints were loaded into R script (Supplementary Data 11). Clustering was performed using the K-means algorithm (dtwclust), grouping genes into eight clusters based on similar temporal dynamic patterns. Euclidean distance was used in clustering to highlight shared trends over time. Mean expression profiles for each cluster were calculated and visualized using ggplot2 to display cluster-specific temporal patterns. Additionally, individual cluster plots were generated using highcharter to provide a dynamic view of each gene’s temporal trajectory within each cluster. The code is openly available on our GitHub repository at https://github.com/Rathalodusk/TimeSeriesClus.

### Statistical information

Statistical analyses for thermal-tolerance and reproductive function experiments were performed using Microsoft Excel or GraphPad. Lifespan and survival assays after heat shock were analyzed using OASIS 2(60). Gene ontology analysis utilized WormCat 2.0(69). Additional statistical analyses, including Fisher’s exact test, were conducted in RStudio. The software and R packages employed for processing sequencing data and performing differential analyses in ATAC-seq and RNA-seq datasets are detailed in the Methods section. All experiments were carried out with at least two biological replicates, with similar results. Details regarding the number of replicates, sample size, types of statistical analyses, p-value cutoffs, and raw data are provided in the corresponding figure legends and/or *source.data*.

## Acknowledgements

Thank you to the Lee Lab members for discussions and suggestions on experimental design and data analysis, and to the lab technicians for support with supplies and preparations. Thank you to the Cornell Genomics Innovation Hub/TREx (Dr. Adrian McNairn) for ATAC-seq advice and to Faraz Ahmed and Dr. Paul Munn for guidance on the analysis pipeline. Thank you to the Cornell Institute of Biotechnology for Illumina sequencing and quality control, with specific appreciation to Dr. Qi Sun and Dr. Jeff Glaubitz for bioinformatics consultation. Thank you to the Cornell Statistical Consulting Unit for statistical advice. Strains were provided by the Caenorhabditis Genetics Center (CGC), and Wormbase.org was used throughout this study(70). This work was funded by NIH AG024425 (to S.S.L.), the Taiwanese Study Abroad Scholarship from Ministry of Education Taiwan (to H.-Y.C.), and the Genomics Innovation Seed Grant from Cornell Institute of Biotechnology.

## Author contributions

H.-Y.C. and S.S.L. designed the study and wrote the paper. S.M. and S.M. conducted partial RNA-seq and ATAC-seq data analyses. C.H., C.M., and S.S. carried out some of the thermal-resistance and RNAi experiments. C.G.D. supervised the analysis pipelines. H.-Y.C. conducted all other experiments and data analyses. S.S.L. secured funding and supervised the experimental plans.

## Declaration of interests

The authors declare no competing interests.

## Declaration of generative AI and AI-assisted technologies

During the preparation of this work the authors used ChatGPT-4o in order to improve language and readibility. After using this tool/service, the authors reviewed and edited the content as needed and take full responsibility for the content of the publication.

## Resource Availability

### Lead contact

Requests for further information and resources should be directed to and will be fulfilled by the lead contact, Siu Sylvia Lee

## Materials availability

This study did not generate new unique reagents.

## Data and code availability

ATAC-seq and RNA-seq raw sequencing data, ATAC-seq BedGraph files for visualization, the RNA-seq raw read count matrix, and the ATAC-seq raw read count matrix for identified ‘consensus peaks’ are in the process of being deposited in NCBI’s Gene Expression Omnibus (GEO) and will be accessible upon request or before the final submission. Details on reference files used for genome alignment (ce11/WBcel235) and other analyses are provided in the STAR Methods section. Raw data for all thermal-resistance, survival, and lifespan experiments; gene lists; peak (associated gene) lists; BED format files of consensus peaks; motif candidate lists; and other data corresponding to the main and supplementary figures can be found in *source.data*. RNAi screening results, gene lists and peak (associated gene) lists in heatmaps from K-means clustering, and GO enrichment lists are available in the Supplementary Information (additional Summplementary files). All custom computer codes used for analyses in this study are available in the Supplementary Information (Supplementary Data 5-10).

## Supporting Information

**Supporting information is listed below, including the following files:**

**File Name: Supplementary Information**

**Description:** Supplementary text and Supplementary Figures 1-8

**File Name: Supplementary Data 1**

**Description:** Gene lists and log2FC values for the indicated comparisons of K-means clusters, originally identified from Morpheus (https://software.broadinstitute.org/morpheus) and further rearranged into a heatmap corresponding to Figure 2c.

**File Name: Supplementary Data 2**

**Description:** Peak (associated-gene) lists and log2FC values for the indicated comparisons of K-means clusters, originally identified from Morpheus (https://software.broadinstitute.org/morpheus) and further rearranged into a heatmap corresponding to Figure 2d.

**File Name: Supplementary Data 3**

**Description:** Gene ontology (GO) analysis summary for the indicated gene lists, identified using WormCat 2.0 (Category 2 and 3)(69), including statistical details, corresponding to Figure 2e-g and Figure 3e-f.

**File Name: Supplementary Data 4**

**Description:** RNAi screening list, including instructions for RNAi feeding time and raw data of individual RNAi gene KD survival results.

**File Name: Supplementary Data 5**

**Description:** RNAi screening list in WT background, including instructions for RNAi feeding time and raw data of individual RNAi gene KD survival results.

**File Name: Supplementary Data 6**

**Description:** ATAC-seq analysis pipeline and istructions for execution in Linux

**File Name: Supplementary Data 7**

**Description:** RNA-seq analysis pipeline and instructions for execution in Linux

**File Name: Supplementary Data 8**

**Description:** R script for ruunning differential expression analysis and generating an MA and Volcano plot.

**File Name: Supplementary Data 9**

**Description:** R script for annotating consensus peaks.

**File Name: Supplementary Data 10**

**Description:** R script for fetching promoter region sequences for motif analysis.

**File Name: Supplementary Data 11**

**Description:** R script for conducting temporal dynamic analysis with time-series clustering and plotting interactive trajectories.

**File Name: Supplementary Data 12Description:** Matrix of raw read counts from RNA-seq data. (Will also be uploaded to GEO metadata.)

**File Name: Supplementary Data 13**

**Description:** Matrix of raw read counts in identified consensus peaks from ATAC-seq data. (Will also be uploaded to GEO metadata.)

**File Name: Supplementary Data 14**

**Description:** Reference Elements for peak annotation, adapted from a published dataset(64).

**File Name: S_source.data**

**Description:** Detailed information and datasets corresponding to the indicated figures in spreadsheet format. Referred to as *source.data* in the main text

## Notes

### Competing Interest Statement

The authors have declared no competing interest.

### Summary of Updates

In the revised manuscript, we incorporated new data and analyses that significantly strengthen the findings and broaden the scope of our study. Specifically, three major updates were made: 1. Integration of multi-omic data from wild-type (WT) worms We included RNA-seq and ATAC-seq datasets from WT worms (Supplementary Fig. 5-7), which were not presented in the previous version. These results reveal that dynamic changes in gene expression and chromatin accessibility across the heat hormesis regimen are highly similar between WT and the germline-less mutant glp-1(ts). This demonstrates that the molecular programs underlying hormetic heat adaptation are largely conserved across genetic backgrounds. 2. Newly identified regulators play roles in heat hormesis in both WT and glp-1(ts) We further tested regulators identified through multi-omic analysis in the glp-1(ts) background and found that mars-1, snpc-4, and fos-1 also enhanced thermal-resistance in WT worms, similar to glp-1(ts). While hsf-1, elt-2, and dpy-27 exhibited context-specific effects (Fig. 4; Supplementary Fig. 7). These findings highlight several putative regulators with conserved roles in heat hormesis across genotypes, while also revealing factors that act in a background-dependent manner. 3. New insights into lifespan extension mechanisms We added new data showing that priming preserves inducibility of heat-shock proteins during early adulthood (Fig. 5d). Together with evidence of transient germline defects, these results suggest that preserved HSP responsiveness, coupled with temporary germline impairment, may underlie the lifespan extension induced by heat hormesis (Fig. 5; Supplementary Fig. 8). In addition to these major additions, we made several technical and presentation improvements to enhance clarity and reproducibility. Figure arrangements were streamlined for narrative coherence, and volcano plots replaced MA plots in Fig. 2a-b and Fig. 3a to better visualize differential RNA expression and chromatin accessibility. Collectively, these revisions expand the scope of the manuscript. The new data provide stronger mechanistic support for both conserved and germline-dependent regulation of heat hormesis, linking chromatin accessibility, gene expression dynamics, and functional outcomes in thermal resistance as well as lifespan.

## References

1. Calabrese EJ. Hormesis: a fundamental concept in biology. Microb Cell [Internet]. 2014 Apr 23 [cited 2025 Feb 16];1(5):145–9. Available from: https://microbialcell.com/researcharticles/hormesis-a-fundamental-concept-in-biology/, https://microbialcell.com/researcharticles/hormesis-a-fundamental-concept-in-biology/

2. Gems D, Partridge L. Stress-Response Hormesis and Aging: “That which Does Not Kill Us Makes Us Stronger.” Cell Metab [Internet]. 2008 Mar 5 [cited 2025 Feb 16];7(3):200–3. Available from: https://www.cell.com/cell-metabolism/abstract/S1550-4131(08)00002-8

3. Mattson MP. Hormesis defined. Ageing Res Rev. 2008 Jan;7(1):1–7.

4. Cypser JR, Johnson TE. Multiple stressors in Caenorhabditis elegans induce stress hormesis and extended longevity. J Gerontol A Biol Sci Med Sci. 2002 Mar;57(3):B109–114.

5. Shama S, Lai CY, Antoniazzi JM, Jiang JC, Jazwinski SM. Heat Stress-Induced Life Span Extension in Yeast. Exp Cell Res [Internet]. 1998 Dec 15 [cited 2025 Feb 16];245(2):379–88. Available from: https://www.sciencedirect.com/science/article/pii/S0014482798942793

6. Yokoyama K, Fukumoto K, Murakami T, Harada S ichi, Hosono R, Wadhwa R, et al. Extended longevity of *Caenorhabditis elegans* by knocking in extra copies of hsp70F, a homolog of mot-2 (mortalin)/mthsp70/Grp75. FEBS Lett [Internet]. 2002 Apr 10 [cited 2025 Feb 16];516(1):53–7. Available from: https://www.sciencedirect.com/science/article/pii/S0014579302024705

7. Cypser JR, Tedesco P, Johnson TE. Hormesis and aging in *Caenorhabditis elegans*. Exp Gerontol [Internet]. 2006 Oct 1 [cited 2025 Feb 16];41(10):935–9. Available from: https://www.sciencedirect.com/science/article/pii/S053155650600283X

8. Le Bourg É, Valenti P, Lucchetta P, Payre F. Effects of mild heat shocks at young age on aging and longevity in Drosophila melanogaster. Biogerontology [Internet]. 2001 Sept 1 [cited 2025 Feb 16];2(3):155–64. Available from: 10.1023/A:1011561107055

9. Hercus MJ, Loeschcke V, Rattan SIS. Lifespan extension of Drosophila melanogaster through hormesis by repeated mild heat stress. Biogerontology. 2003;4(3):149–56.

10. Rattan SIS. Hormetic Modulation of Aging and Longevity by Mild Heat Stress. Dose-Response [Internet]. 2005 Oct 1 [cited 2025 Feb 16];3(4):dose-response.003.04.008. Available from: 10.2203/dose-response.003.04.008

11. Rattan SIS, Fernandes RA, Demirovic D, Dymek B, Lima CF. Heat Stress and Hormetin-Induced Hormesis in Human Cells: Effects on Aging, Wound Healing, Angiogenesis, and Differentiation. Dose-Response [Internet]. 2008 Sept 10 [cited 2025 Feb 16];7(1):90–103. Available from: https://www.ncbi.nlm.nih.gov/pmc/articles/PMC2664638/

12. Mane NR, Gajare KA, Deshmukh AA. Mild heat stress induces hormetic effects in protecting the primary culture of mouse prefrontal cerebrocortical neurons from neuropathological alterations. IBRO Rep [Internet]. 2018 Dec 1 [cited 2025 Feb 16];5:110–5. Available from: https://www.sciencedirect.com/science/article/pii/S2451830118300724

13. Verbeke P, Deries M, Clark BFC, Rattan SIS. Hormetic action of mild heat stress decreases the inducibility of protein oxidation and glycoxidation in human fibroblasts. Biogerontology. 2002;3(1– 2):117–20.

14. Laukkanen T, Kunutsor SK, Khan H, Willeit P, Zaccardi F, Laukkanen JA. Sauna bathing is associated with reduced cardiovascular mortality and improves risk prediction in men and women: a prospective cohort study. BMC Med [Internet]. 2018 Nov 29 [cited 2025 Feb 16];16(1):219. Available from: 10.1186/s12916-018-1198-0

15. Patrick RP, Johnson TL. Sauna use as a lifestyle practice to extend healthspan. Exp Gerontol [Internet]. 2021 Oct 15 [cited 2025 Feb 16];154:111509. Available from: https://www.sciencedirect.com/science/article/pii/S0531556521002916

16. Lithgow GJ, White TM, Melov S, Johnson TE. Thermotolerance and extended life-span conferred by single-gene mutations and induced by thermal stress. Proc Natl Acad Sci U S A [Internet]. 1995 Aug 1 [cited 2025 Feb 16];92(16):7540–4. Available from: https://www.ncbi.nlm.nih.gov/pmc/articles/PMC41375/

17. McColl G, Rogers AN, Alavez S, Hubbard AE, Melov S, Link CD, et al. Insulin-like Signaling Determines Survival During Stress via Post Transcriptional Mechanisms in C. elegans. Cell Metab [Internet]. 2010 Sept 8 [cited 2025 Feb 16];12(3):260–72. Available from: https://www.ncbi.nlm.nih.gov/pmc/articles/PMC2945254/

18. Butov A, Johnson T, Cypser J, Sannikov I, Volkov M, Sehl M, et al. Hormesis and debilitation effects in stress experiments using the nematode worm *Caenorhabditis elegans*: the model of balance between cell damage and HSP levels. Exp Gerontol [Internet]. 2001 Dec 1 [cited 2025 Feb 16];37(1):57–66. Available from: https://www.sciencedirect.com/science/article/pii/S0531556501001619

19. Kumsta C, Chang JT, Schmalz J, Hansen M. Hormetic heat stress and HSF-1 induce autophagy to improve survival and proteostasis in C. elegans. Nat Commun [Internet]. 2017 Feb 15 [cited 2025 Feb 16];8(1):14337. Available from: https://www.nature.com/articles/ncomms14337

20. Zhang B, Xiao R, Ronan EA, He Y, Hsu AL, Liu J, et al. Environmental Temperature Differentially Modulates *C. elegans* Longevity through a Thermosensitive TRP Channel. Cell Rep [Internet]. 2015 June 9 [cited 2025 Mar 6];11(9):1414–24. Available from: https://www.sciencedirect.com/science/article/pii/S2211124715004982

21. Zhou L, He B, Deng J, Pang S, Tang H. Histone acetylation promotes long-lasting defense responses and longevity following early life heat stress. PLOS Genet [Internet]. 2019 Apr 29 [cited 2025 Feb 16];15(4):e1008122. Available from: https://journals.plos.org/plosgenetics/article?id=10.1371/journal.pgen.1008122

22. Kourtis N, Nikoletopoulou V, Tavernarakis N. Small heat-shock proteins protect from heat-stroke-associated neurodegeneration. Nature [Internet]. 2012 Oct [cited 2025 Mar 5];490(7419):213–8. Available from: https://www.nature.com/articles/nature11417

23. Wang S, You M, Wang C, Zhang Y, Fan C, Yan S. Heat shock pretreatment induced cadmium resistance in the nematode *Caenorhabditis elegans* is depend on transcription factors DAF-16 and HSF-1. Environ Pollut [Internet]. 2020 June 1 [cited 2025 Mar 5];261:114081. Available from: https://www.sciencedirect.com/science/article/pii/S0269749119357008

24. Wan QL, Meng X, Dai W, Luo Z, Wang C, Fu X, et al. N6-methyldeoxyadenine and histone methylation mediate transgenerational survival advantages induced by hormetic heat stress. Sci Adv [Internet]. 2021 Jan [cited 2025 Mar 10];7(1):eabc3026. Available from: https://www.science.org/doi/10.1126/sciadv.abc3026

25. Huang J, Wang K, Wang M, Wu Z, Xie G, Peng Y, et al. One-day thermal regime extends the lifespan in Caenorhabditis elegans. Comput Struct Biotechnol J [Internet]. 2023 Jan 1 [cited 2025 Mar 22];21:495–505. Available from: https://www.sciencedirect.com/science/article/pii/S2001037022005748

26. Treinin M, Shliar J, Jiang H, Powell-Coffman JA, Bromberg Z, Horowitz M. HIF-1 is required for heat acclimation in the nematode Caenorhabditis elegans. Physiol Genomics. 2003 June 24;14(1):17–24.

27. Cypser JR, Johnson TE. Hormesis in Caenorhabditis elegans dauer-defective mutants. Biogerontology. 2003;4(4):203–14.

28. Jiang WI, De Belly H, Wang B, Wong A, Kim M, Oh F, et al. Early-life stress triggers long-lasting organismal resilience and longevity via tetraspanin. Sci Adv [Internet]. 2024 Jan 24 [cited 2025 Feb 16];10(4):eadj3880. Available from: https://www.science.org/doi/full/10.1126/sciadv.adj3880

29. Xu F, Li R, von Gromoff ED, Drepper F, Knapp B, Warscheid B, et al. Reprogramming of the transcriptome after heat stress mediates heat hormesis in Caenorhabditis elegans. Nat Commun [Internet]. 2023 July 13 [cited 2025 Feb 16];14(1):4176. Available from: https://www.nature.com/articles/s41467-023-39882-8

30. Schreiner WP, Pagliuso DC, Garrigues JM, Chen JS, Aalto AP, Pasquinelli AE. Remodeling of the Caenorhabditis elegans non-coding RNA transcriptome by heat shock. Nucleic Acids Res [Internet]. 2019 Oct 10 [cited 2025 Feb 16];47(18):9829–41. Available from: 10.1093/nar/gkz693

31. Kim Y, Sun H. Functional genomic approach to identify novel genes involved in the regulation of oxidative stress resistance and animal lifespan. Aging Cell. 2007 Aug;6(4):489–503.

32. Shi Y, Mosser DD, Morimoto RI. Molecularchaperones as HSF1-specific transcriptional repressors. Genes Dev [Internet]. 1998 Mar 1 [cited 2025 Feb 16];12(5):654–66. Available from: http://genesdev.cshlp.org/content/12/5/654

33. Yang W, Dierking K, Schulenburg H. WormExp: a web-based application for a Caenorhabditis elegans-specific gene expression enrichment analysis. Bioinformatics [Internet]. 2016 Mar 15 [cited 2025 Sept 12];32(6):943–5. Available from: 10.1093/bioinformatics/btv667

34. Labbadia J, Morimoto RI. Repression of the Heat Shock Response Is a Programmed Event at the Onset of Reproduction. Mol Cell. 2015 Aug 20;59(4):639–50.

35. Calabrese EJ, Bachmann KA, Bailer AJ, Bolger PM, Borak J, Cai L, et al. Biological stress response terminology: Integrating the concepts of adaptive response and preconditioning stress within a hormetic dose–response framework. Toxicol Appl Pharmacol [Internet]. 2007 July 1 [cited 2025 Feb 16];222(1):122–8. Available from: https://www.sciencedirect.com/science/article/pii/S0041008X07001032

36. Starks RR, Biswas A, Jain A, Tuteja G. Combined analysis of dissimilar promoter accessibility and gene expression profiles identifies tissue-specific genes and actively repressed networks. Epigenetics Chromatin [Internet]. 2019 Feb 22 [cited 2025 Feb 16];12(1):16. Available from: 10.1186/s13072-019-0260-2

37. Morimoto RI. The heat shock response: systems biology of proteotoxic stress in aging and disease. Cold Spring Harb Symp Quant Biol. 2011;76:91–9.

38. Lindquist S, Craig EA. THE HEAT-SHOCK PROTEINS. Annu Rev Genet [Internet]. 1988 Dec 1 [cited 2025 Feb 16];22(Volume 22, 1988):631–77. Available from: https://www.annualreviews.org/content/journals/10.1146/annurev.ge.22.120188.003215

39. McGhee JD, Fukushige T, Krause MW, Minnema SE, Goszczynski B, Gaudet J, et al. ELT-2 is the predominant transcription factor controlling differentiation and function of the C. elegans intestine, from embryo to adult. Dev Biol. 2009 Mar 15;327(2):551–65.

40. Wiesenfahrt T, Berg JY, Osborne Nishimura E, Robinson AG, Goszczynski B, Lieb JD, et al. The function and regulation of the GATA factor ELT-2 in the C. elegans endoderm. Development [Internet]. 2016 Feb 1 [cited 2025 Feb 16];143(3):483–91. Available from: 10.1242/dev.130914

41. Kerry S, TeKippe M, Gaddis NC, Aballay A. GATA Transcription Factor Required for Immunity to Bacterial and Fungal Pathogens. PLOS ONE [Internet]. 2006 Dec 20 [cited 2025 Feb 16];1(1):e77. Available from: https://journals.plos.org/plosone/article?id=10.1371/journal.pone.0000077

42. Kovács D, Biró JB, Ahmed S, Kovács M, Sigmond T, Hotzi B, et al. Age-dependent heat shock hormesis to HSF-1 deficiency suggests a compensatory mechanism mediated by the unfolded protein response and innate immunity in young Caenorhabditis elegans. Aging Cell [Internet]. 2024 [cited 2025 Feb 16];23(10):e14246. Available from: https://onlinelibrary.wiley.com/doi/abs/10.1111/acel.14246

43. Meyer BJ. The X chromosome in *C. elegans* sex determination and dosage compensation. Curr Opin Genet Dev [Internet]. 2022 June 1 [cited 2025 Mar 7];74:101912. Available from: https://www.sciencedirect.com/science/article/pii/S0959437X22000211

44. Anderson EC, Frankino PA, Higuchi-Sanabria R, Yang Q, Bian Q, Podshivalova K, et al. X Chromosome Domain Architecture Regulates Caenorhabditis elegans Lifespan but Not Dosage Compensation. Dev Cell [Internet]. 2019 Oct 21 [cited 2025 Mar 7];51(2):192-207.e6. Available from: https://www.cell.com/developmental-cell/abstract/S1534-5807(19)30664-1

45. Shaulian E, Karin M. AP-1 as a regulator of cell life and death. Nat Cell Biol [Internet]. 2002 May [cited 2025 Feb 16];4(5):E131–6. Available from: https://www.nature.com/articles/ncb0502-e131

46. Zhang Z, Liu L, Twumasi-Boateng K, Block DHS, Shapira M. FOS-1 functions as a transcriptional activator downstream of the C. elegans JNK homolog KGB-1. Cell Signal. 2017 Jan;30:1–8.

47. Gerke P, Keshet A, Mertenskötter A, Paul RJ. The JNK-Like MAPK KGB-1 of Caenorhabditis Elegans Promotes Reproduction, Lifespan, and Gene Expressions for Protein Biosynthesis and Germline Homeostasis but Interferes with Hyperosmotic Stress Tolerance. Cell Physiol Biochem [Internet]. 2014 Nov 25 [cited 2025 Feb 16];34(6):1951–73. Available from: 10.1159/000366392

48. Lynch CJ, Richart L, Serrano M. A pattern emerges in chromatin aging: AP-1 steals the show. Cell Metab [Internet]. 2024 Aug 6 [cited 2025 Feb 16];36(8):1639–41. Available from: https://www.cell.com/cell-metabolism/abstract/S1550-4131(24)00286-9

49. Patrick R, Naval-Sanchez M, Deshpande N, Huang Y, Zhang J, Chen X, et al. The activity of early-life gene regulatory elements is hijacked in aging through pervasive AP-1-linked chromatin opening. Cell Metab [Internet]. 2024 Aug 6 [cited 2025 Feb 16];36(8):1858-1881.e23. Available from: https://www.cell.com/cell-metabolism/abstract/S1550-4131(24)00231-6

50. Kasper DM, Wang G, Gardner KE, Johnstone TG, Reinke V. The C. elegans SNAPc Component SNPC-4 Coats piRNA Domains and Is Globally Required for piRNA Abundance. Dev Cell [Internet]. 2014 Oct 27 [cited 2025 Feb 16];31(2):145–58. Available from: https://www.cell.com/developmental-cell/abstract/S1534-5807(14)00621-2

51. Weng C, Kosalka J, Berkyurek AC, Stempor P, Feng X, Mao H, et al. The USTC co-opts an ancient machinery to drive piRNA transcription in C. elegans. Genes Dev. 2019 Jan 1;33(1–2):90–102.

52. Belicard T, Jareosettasin P, Sarkies P. The piRNA pathway responds to environmental signals to establish intergenerational adaptation to stress. BMC Biol [Internet]. 2018 Sept 18 [cited 2025 Feb 16];16(1):103. Available from: 10.1186/s12915-018-0571-y

53. Batista PJ, Ruby JG, Claycomb JM, Chiang R, Fahlgren N, Kasschau KD, et al. PRG-1 and 21U-RNAs interact to form the piRNA complex required for fertility in C. elegans. Mol Cell. 2008 July 11;31(1):67–78.

54. Wang A, Song Z, Zhang X, Xiao L, Feng Y, Qi C, et al. MARS1 mutations linked to familial trigeminal neuralgia via the integrated stress response. J Headache Pain [Internet]. 2023 Jan 14 [cited 2025 Feb 16];24(1):4. Available from: https://www.ncbi.nlm.nih.gov/pmc/articles/PMC9840295/

55. Arantes-Oliveira N, Apfeld J, Dillin A, Kenyon C. Regulation of Life-Span by Germ-Line Stem Cells in Caenorhabditis elegans. Science [Internet]. 2002 Jan 18 [cited 2025 Oct 6];295(5554):502–5. Available from: https://www.science.org/doi/full/10.1126/science.1065768

56. Moses E, Atlan T, Sun X, Franěk R, Siddiqui A, Marinov GK, et al. The killifish germline regulates longevity and somatic repair in a sex-specific manner. Nat Aging. 2024 June;4(6):791–813.

57. Flatt T, Min KJ, D’Alterio C, Villa-Cuesta E, Cumbers J, Lehmann R, et al. Drosophila germ-line modulation of insulin signaling and lifespan. Proc Natl Acad Sci [Internet]. 2008 Apr 29 [cited 2025 Oct 6];105(17):6368–73. Available from: https://www.pnas.org/doi/10.1073/pnas.0709128105

58. Hsin H, Kenyon C. Signals from the reproductive system regulate the lifespan of C. elegans. Nature. 1999 May 27;399(6734):362–6.

59. Bazopoulou D, Knoefler D, Zheng Y, Ulrich K, Oleson BJ, Xie L, et al. Developmental ROS individualizes organismal stress resistance and lifespan. Nature [Internet]. 2019 Dec [cited 2025 Feb 16];576(7786):301–5. Available from: https://www.nature.com/articles/s41586-019-1814-y

60. Han SK, Lee D, Lee H, Kim D, Son HG, Yang JS, et al. OASIS 2: online application for survival analysis 2 with features for the analysis of maximal lifespan and healthspan in aging research. Oncotarget. 2016 Aug 30;7(35):56147–52.

61. Kamath RS, Ahringer J. Genome-wide RNAi screening in *Caenorhabditis elegans*. Methods [Internet]. 2003 Aug 1 [cited 2025 Feb 16];30(4):313–21. Available from: https://www.sciencedirect.com/science/article/pii/S1046202303000501

62. Rual JF, Ceron J, Koreth J, Hao T, Nicot AS, Hirozane-Kishikawa T, et al. Toward Improving Caenorhabditis elegans Phenome Mapping With an ORFeome-Based RNAi Library. Genome Res [Internet]. 2004 Oct [cited 2025 Feb 16];14(10b):2162–8. Available from: https://www.ncbi.nlm.nih.gov/pmc/articles/PMC528933/

63. Daugherty AC, Yeo RW, Buenrostro JD, Greenleaf WJ, Kundaje A, Brunet A. Chromatin accessibility dynamics reveal novel functional enhancers in C. elegans. Genome Res. 2017 Dec;27(12):2096–107.

64. Chen RAJ, Down TA, Stempor P, Chen QB, Egelhofer TA, Hillier LW, et al. The landscape of RNA polymerase II transcription initiation in C. elegans reveals promoter and enhancer architectures. Genome Res. 2013 Aug;23(8):1339–47.

65. Jänes J, Dong Y, Schoof M, Serizay J, Appert A, Cerrato C, et al. Chromatin accessibility dynamics across C. elegans development and ageing. Lee SS, Tyler JK, editors. eLife [Internet]. 2018 Oct 26 [cited 2025 Feb 16];7:e37344. Available from: 10.7554/eLife.37344

66. Bailey TL, Johnson J, Grant CE, Noble WS. The MEME Suite. Nucleic Acids Res [Internet]. 2015 July 1 [cited 2025 Feb 18];43(W1):W39–49. Available from: 10.1093/nar/gkv416

67. Rauluseviciute I, Riudavets-Puig R, Blanc-Mathieu R, Castro-Mondragon JA, Ferenc K, Kumar V, et al. JASPAR 2024: 20th anniversary of the open-access database of transcription factor binding profiles. Nucleic Acids Res [Internet]. 2024 Jan 5 [cited 2025 Feb 18];52(D1):D174–82. Available from: 10.1093/nar/gkad1059

68. Weirauch MT, Yang A, Albu M, Cote AG, Montenegro-Montero A, Drewe P, et al. Determination and inference of eukaryotic transcription factor sequence specificity. Cell. 2014 Sept 11;158(6):1431–43.

69. Holdorf AD, Higgins DP, Hart AC, Boag PR, Pazour GJ, Walhout AJM, et al. WormCat: An Online Tool for Annotation and Visualization of Caenorhabditis elegans Genome-Scale Data. Genetics [Internet]. 2020 Feb 1 [cited 2025 Feb 16];214(2):279–94. Available from: 10.1534/genetics.119.302919

70. Davis P, Zarowiecki M, Arnaboldi V, Becerra A, Cain S, Chan J, et al. WormBase in 2022—data, processes, and tools for analyzing Caenorhabditis elegans. Genetics [Internet]. 2022 Feb 4 [cited 2025 Feb 17];220(4):iyac003. Available from: https://www.ncbi.nlm.nih.gov/pmc/articles/PMC8982018/

