## Supplementary Information for "Transcriptomic and chromatin accessibility profiling unveils new regulators of heat hormesis in *Caenorhabditis elegans*"

- Supplementary text
- Supplementary Figures 1-8
- Descriptions of additional supplementary files

### Supplementary text

#### RNAseq and ATACseq data are of high quality

The data demonstrated good correlation among biological replicates for both RNA-seq and ATAC-seq results (Supplementary Fig. 2a-j), indicating high reproducibility of the experiments. Additionally, as expected(1), the ATAC-seq signals were enriched at regions surrounding transcriptional start sites (TSS)(2) (Supplementary Fig. 2l), again supporting the high quality of the data (*source.data*: RNA-seq QC, ATAC-seq QC). For RNA-seq, we detected expression of approximately 13,000 genes among the different experimental conditions. For ATAC-seq, we identified a total of 30,404 consensus peaks (*source.data*). In accordance with published analysis pipeline(3,4), we employed the narrow peak calling tool (-f BAMPE --bdg --SPMR --gsize ce -q 0.05 --call-summits) of MACS2 and the peaks were further subdivided based on their summits to achieve optimal resolution of consensus peaks. These peaks were additionally associated with 19,352 genes based on published dataset(4) (see Methods for detail).

#### Temporal trajectories of RNA expression change between primed and naive worms

To visualize the temporal trajectoris of the RNA expression differences between primed and naive worms, we conducted clustering analysis that integrated the gene expression differences between the two groups across the three timepoints. This analysis categorized the genes exhibiting significant RNA expression differences between primed and naive groups into eight distinct trajectories (Supplementary Fig. 4c; *source.data*). The distinct trajectories highlighted different degrees of RNA expression differences between primed and naive groups in response to priming and subsequent HS. Most of the trajectories (7 out of 8) followed a general pattern of peaking immediately after priming (timepoint 1) and returning to baseline by the recovery phase (timepoint 2), consistent with our earlier conclusion that the majority of the priming induced RNA expression changes are restored upon a 12-hour recovery (Fig. 2a, c: C.1). Three trajectories (2, 6, 7) represented genes exhibiting differential RNA expression between primed and naive group upon a 6-hour HS (timepoint 3). Clusters 2 and 7, representing genes associated with lower RNA expression in primed compared to naive worms, are enriched for the GO terms detoxification, pathogen stress response, and lipid metabolism, aligning with our previous findings (Supplementary Fig. 4d). Cluster 2 captured genes that showed no significant differences between primed and naive at timepoint 1 but exhibited substantial differences at timepoint 3. On the other hand, clusters 7 and 6 captured the relatively small number of genes that showed persistently lower or higher RNA expression respectfully in primed compared to naive worms across the heat hormesis regimen. Therefore, this time-series analysis led us to a similar conclusion that most priming-induced gene expression changes are transient, but also highlighted different waves of RNA expression differences between the primed and naive groups, some of which likely contribute to the stress resilience of primed worms.

We additionally compared the fold change in RNA expression or chromatin accessibility after HS in the primed vs naive worms and observed a high degree of concordance (Supplementary Fig. 4b), indicating that there was no global shift in HS-induced changes in RNA expression and chromatin accessibility between the primed and naive groups.

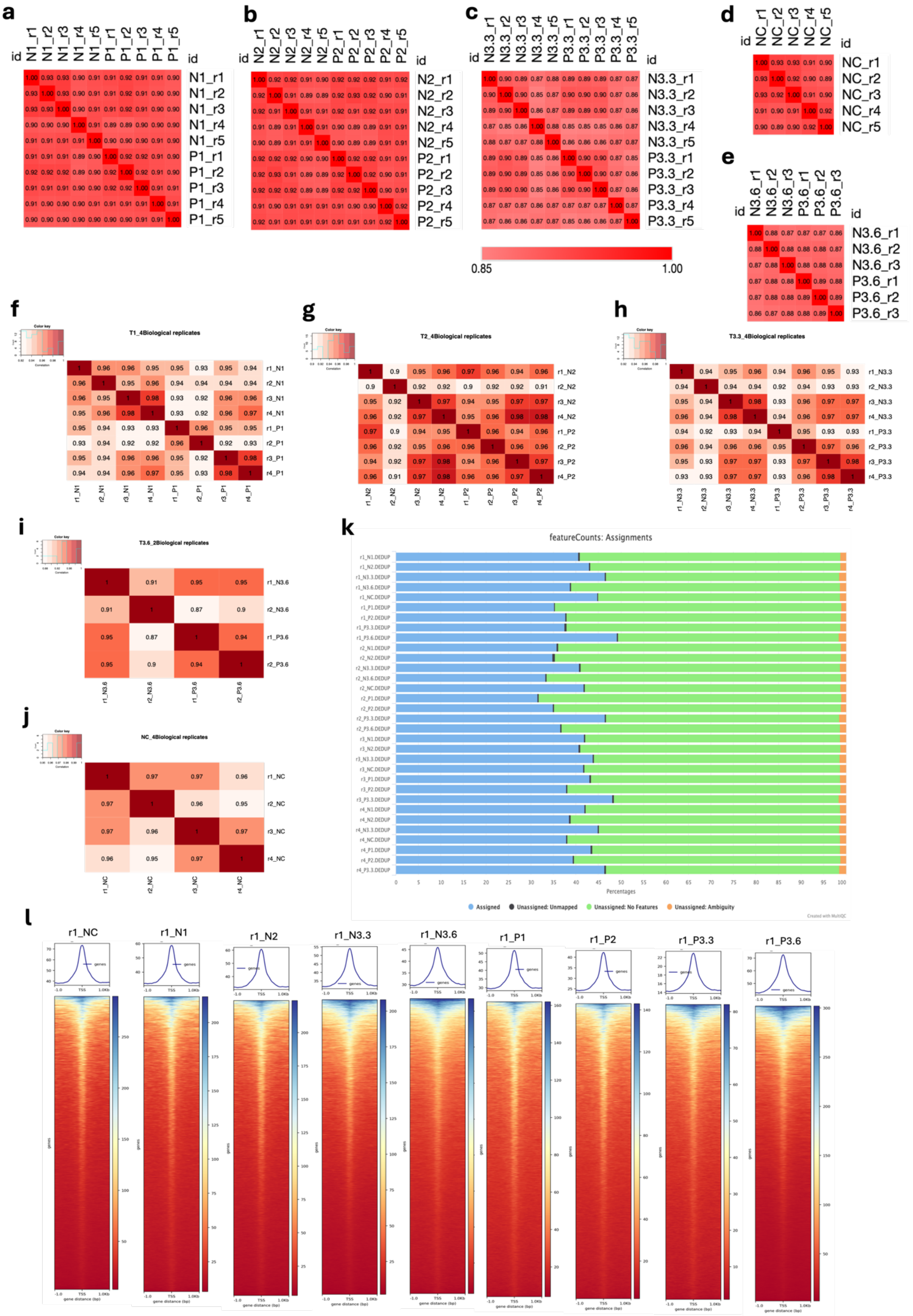
a
b
c
d
e
f
T1_4Biological replicates
g
T2_4Biological replicates
h
T3.3_4Biological replicates
i
T3.6_2Biological replicates
j
NC_4Biological replicates
k
featureCounts: Assignments
Percentages
Assigned
Unassigned: Unmapped
Unassigned: No Features
Unassigned: Ambiguity
Created with MultiQC
l
r1_NC
r1_N1
r1_N2
r1_N3.3
r1_N3.6
r1_P1
r1_P2
r1_P3.3
r1_P3.6

**Supplementary Figure 1: Quality control for transcriptomic and chromatin accessibility profiles in *glp-1(ts)*.**

Spearman's correlation analysis of RNA-seq profiles for both naive and primed groups of *glp-1(ts)* at timepoints 1 (**a**), 2 (**b**), 3.3 (**c**), 3.6 (**e**) and negative control (**d**) across independent replicates. Pearson's correlation analysis of ATAC-seq profiles for both naive and primed groups at timepoints 1 (**f**), 2 (**g**), 3.3 (**h**), 3.6 (**i**), and negative control (**j**) across independent replicates. **(k)** Fraction of Reads in Peaks (FRiP) scores for individual samples, calculated using MultiQC based on featureCounts. 'Assigned featureCounts' indicates mapped reads counted within identified consensus peaks (*source.data*). 'Unassigned: no Features' indicates mapped reads not counted in consensus peaks. The FRiP scores within the identified consensus peaks ranged from 31% to 49% across all samples, affirming the good quality of the data. Furthermore, the FRiP scores among biological replicates were highly consistent, further supporting the reproducibility of our datasets. **(l)** TSS enrichment for all experimental groups in a representative biological replicate (r1): The top panel displays profile plots aggregating read coverage within 1kb upstream and downstream around TSS for all genes across the genome. The bottom panel displays heatmaps showing individual gene coverage, with each row corresponding to the TSS of a single gene, extending 1kb upstream and downstream. Colors in the heatmap indicating the level of read coverage. The plots for the remaining replicates can be found in the *source.data*.

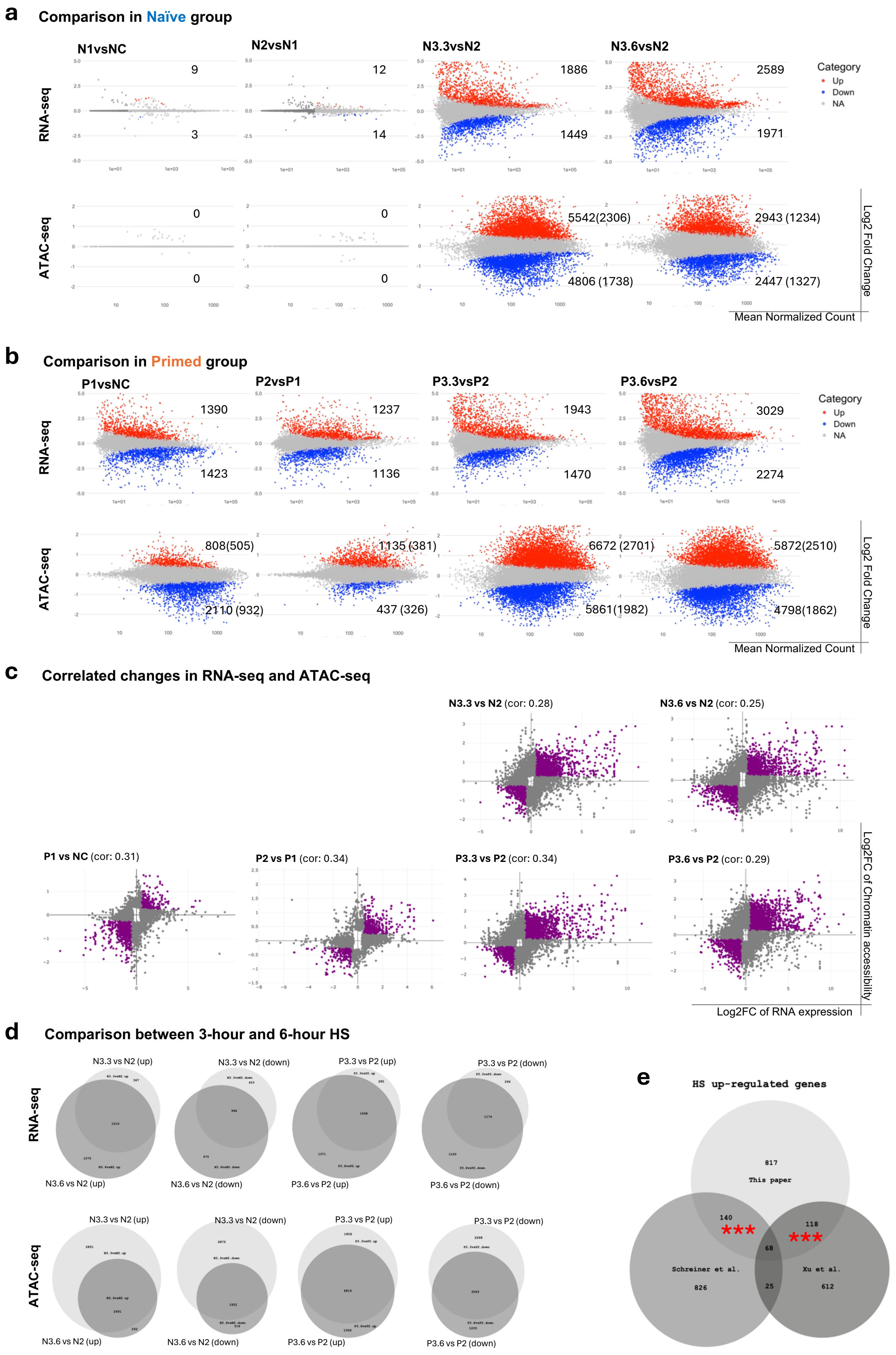
a
Comparison in Naïve group
N1vsNC
N2vsN1
N3.3vsN2
N3.6vsN2
RNA-seq
9
3
12
14
1886
1449
2589
1971
Category
Up
Down
NA
ATAC-seq
0
0
0
0
5542(2306)
4806 (1738)
2943 (1234)
2447 (1327)
Log2 Fold Change
Mean Normalized Count
b
Comparison in Primed group
P1vsNC
P2vsP1
P3.3vsP2
P3.6vsP2
RNA-seq
1390
1423
1237
1136
1943
1470
3029
2274
Category
Up
Down
NA
ATAC-seq
808(505)
2110 (932)
1135 (381)
437 (326)
6672 (2701)
5861(1982)
5872(2510)
4798(1862)
Log2 Fold Change
Mean Normalized Count
c
Correlated changes in RNA-seq and ATAC-seq
N3.3 vs N2 (cor: 0.28)
N3.6 vs N2 (cor: 0.25)
P1 vs NC (cor: 0.31)
P2 vs P1 (cor: 0.34)
P3.3 vs P2 (cor: 0.34)
P3.6 vs P2 (cor: 0.29)
Log2FC of Chromatin accessibility
Log2FC of RNA expression
d
Comparison between 3-hour and 6-hour HS
RNA-seq
N3.3 vs N2 (up)
N3.3 vs N2 (down)
P3.3 vs P2 (up)
P3.3 vs P2 (down)
N3.6 vs N2 (up)
N3.6 vs N2 (down)
P3.6 vs P2 (up)
P3.6 vs P2 (down)
ATAC-seq
N3.3 vs N2 (up)
N3.3 vs N2 (down)
P3.3 vs P2 (up)
P3.3 vs P2 (down)
N3.6 vs N2 (up)
N3.6 vs N2 (down)
P3.6 vs P2 (up)
P3.6 vs P2 (down)
e
HS up-regulated genes
817
This paper
140
118
68
Schreiner et al.
826
25
Xu et al.
612

**Supplementary Figure 2: Differential analyses across timepoints and correlation between RNA expression and chromatin accessibility in *glp-1(ts)*.**

MA plots display log2FC of gene expression from RNA-seq (top panel) and chromatin accessibility from ATAC-seq (bottom panel) for the indicated comparison in naive **(a)** and primed **(b)** groups. Differential analyses were calculated using DESeq2. Significant changes (p-adj < 0.05) are marked in red for upregulation (log2FC > 0) and in blue for downregulation (log2FC < 0), while unchanged are marked in grey (p-adj >= 0.05). Numbers indicate the count of significant genes for RNA-seq data and significant peaks (and their associated genes in brackets) for ATAC-seq data for each plot. **(c)** Scatter plots display genes with significant changes identified in either RNA expression or chromatin accessibility for the indicated comparisons. Genes with correlated changes between RNA-seq and ATAC-seq data are highlighted in purple based on defined filter criteria: log2FC RNA expression > 0.5, < -0.5; log2FC Chromatin accessibility > 0.25, < -0.25. These genes showed upregulation in both RNA expression and Chromatin accessibility or downregulation in both RNA expression and Chromatin accessibility. Genes without correlated changes are in grey. (**d)** Venn diagrams display the number of significantly differentially expressed genes (top panel) or peaks (bottom panel) for the 3-hour HS (light grey) and their overlap with the 6-hour HS (dark grey) in the indicated comparison. Substantial overlaps suggest that 35°C heat shock for 3 or 6 hours elicited many similar changes in RNA expression and chromatin accessibility. **(e)** Venn diagrams display the number of heat shock/stress (HS)-induced genes identified from RNA-seq, comparing our data (N3.3vsN2, Supplementary Fig. 3a) with two published datasets (Schreiner et al. and Xu et al.). Details of experimental setup and HS conditions can be found in *souce.data*. Fisher exact tests were conducted to determine if the overlap between this study (This paper) and published datasets are statistically significant. *** Indicates p < 2.2e-16.

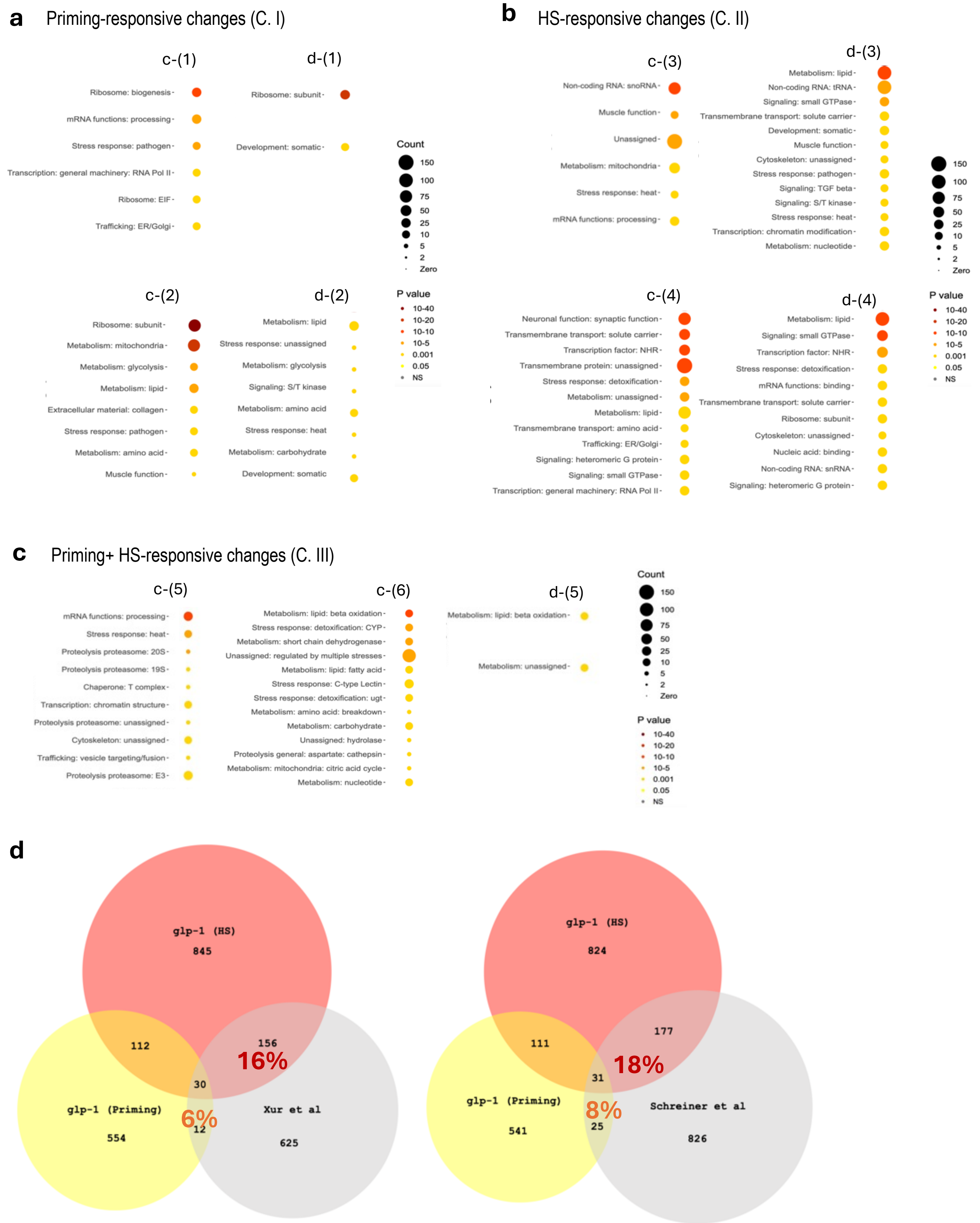
a
Priming-responsive changes (C. I)
c-(1)
Ribosome: biogenesis
mRNA functions: processing
Stress response: pathogen
Transcription: general machinery: RNA Pol II
Ribosome: EIF
Trafficking: ER/Golgi
d-(1)
Ribosome: subunit
Development: somatic
Count
150
100
75
50
25
10
5
2
Zero
c-(2)
Ribosome: subunit
Metabolism: mitochondria
Metabolism: glycolysis
Metabolism: lipid
Extracellular material: collagen
Stress response: pathogen
Metabolism: amino acid
Muscle function
d-(2)
Metabolism: lipid
Stress response: unassigned
Metabolism: glycolysis
Signaling: S/T kinase
Metabolism: amino acid
Stress response: heat
Metabolism: carbohydrate
Development: somatic
P value
10-40
10-20
10-10
10-5
0.001
0.05
NS
b
HS-responsive changes (C. II)
c-(3)
Non-coding RNA: snoRNA
Muscle function
Unassigned
Metabolism: mitochondria
Stress response: heat
mRNA functions: processing
d-(3)
Metabolism: lipid
Non-coding RNA: tRNA
Signaling: small GTPase
Transmembrane transport: solute carrier
Development: somatic
Muscle function
Cytoskeleton: unassigned
Stress response: pathogen
Signaling: TGF beta
Signaling: S/T kinase
Stress response: heat
Transcription: chromatin modification
Metabolism: nucleotide
c-(4)
Neuronal function: synaptic function
Transmembrane transport: solute carrier
Transcription factor: NHR
Transmembrane protein: unassigned
Stress response: detoxification
Metabolism: unassigned
Metabolism: lipid
Transmembrane transport: amino acid
Trafficking: ER/Golgi
Signaling: heteromeric G protein
Signaling: small GTPase
Transcription: general machinery: RNA Pol II
d-(4)
Metabolism: lipid
Signaling: small GTPase
Transcription factor: NHR
Stress response: detoxification
mRNA functions: binding
Transmembrane transport: solute carrier
Ribosome: subunit
Cytoskeleton: unassigned
Nucleic acid: binding
Non-coding RNA: snRNA
Signaling: heteromeric G protein
c
Priming+ HS-responsive changes (C. III)
c-(5)
mRNA functions: processing
Stress response: heat
Proteolysis proteasome: 20S
Proteolysis proteasome: 19S
Chaperone: T complex
Transcription: chromatin structure
Proteolysis proteasome: unassigned
Cytoskeleton: unassigned
Trafficking: vesicle targeting/fusion
Proteolysis proteasome: E3
c-(6)
Metabolism: lipid: beta oxidation
Stress response: detoxification: CYP
Metabolism: short chain dehydrogenase
Unassigned: regulated by multiple stresses
Metabolism: lipid: fatty acid
Stress response: C-type Lectin
Stress response: detoxification: ugt
Metabolism: amino acid: breakdown
Metabolism: carbohydrate
Unassigned: hydrolase
Proteolysis general: aspartate: cathepsin
Metabolism: mitochondria: citric acid cycle
Metabolism: nucleotide
d-(5)
Metabolism: lipid: beta oxidation
Metabolism: unassigned
d
glp-1 (HS)
845
112
156
16%
30
glp-1 (Priming)
6%
12
Xur et al
554
625
glp-1 (HS)
824
111
177
18%
31
glp-1 (Priming)
8%
25
Schreiner et al
541
826

**Supplementary Figure 3: Gene Ontology signatures and comparison with published heat-stress datasets.**

Wormcat GO enrichment analysis for genes in C. I **(a)** and C. II **(b),** C. III **(c)** (Fig. 2c-d). Wormcat p-values are determined by one-side Fisher test with FDR correction. Gene and peak lists, and Wormcat outputs, are provided in the Supplementary Data 3. **(d)** Venn diagrams showing overlaps between priming-induced upregulated genes (yellow) and HS-induced upregulated genes (red) identified in *glp-1(ts)* in this study, and HS-upregulated genes (gray) from published datasets by Xu et al. (left) and Schreiner et al. (right). Percentages indicate the proportion of shared HS-upregulated genes between studies. Experimental conditions are indicated below: **glp-1 (Priming):** *glp-1(ts)*, day 2 adults, 30 °C for 6hr vs. 20 °C (P1 vs NC). **glp-1 (HS):** *glp-1(ts)*, day 2 + 12 h adults, 35 °C for 3hr vs. 20 °C (N3.3 vs N2). **Xu et al.:** N2, day 1 adults, 35 °C for 1 h vs. 20 °C. **Schreiner et al.:** N2, L4 larvae, 35 °C for 4 h vs. 20 °C.

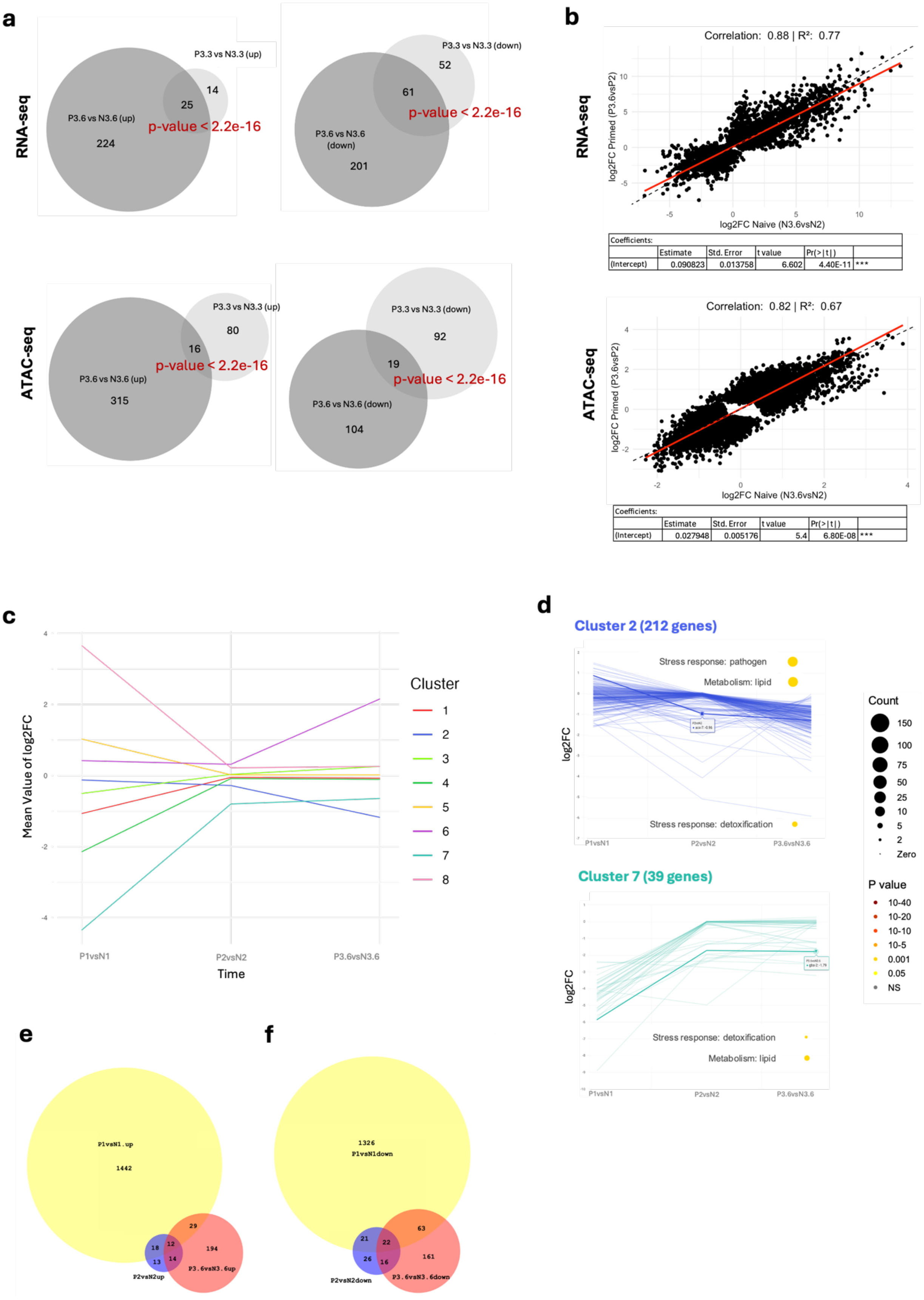
a
RNA-seq
P3.3 vs N3.3 (up)
P3.6 vs N3.6 (up)
224
25
14
p-value < 2.2e-16
P3.3 vs N3.3 (down)
P3.6 vs N3.6 (down)
201
61
52
p-value < 2.2e-16
ATAC-seq
P3.3 vs N3.3 (up)
P3.6 vs N3.6 (up)
315
16
80
p-value < 2.2e-16
P3.3 vs N3.3 (down)
P3.6 vs N3.6 (down)
104
19
92
p-value < 2.2e-16
b
RNA-seq
Correlation: 0.88 | R²: 0.77
log2FC Primed (P3.6vsP2)
log2FC Naive (N3.6vsN2)
Coefficients:
Estimate | Std. Error | t value | Pr(>|t|)
(Intercept) | 0.090823 | 0.013758 | 6.602 | 4.40E-11 | ***
ATAC-seq
Correlation: 0.82 | R²: 0.67
log2FC Primed (P3.6vsP2)
log2FC Naive (N3.6vsN2)
Coefficients:
Estimate | Std. Error | t value | Pr(>|t|)
(Intercept) | 0.027948 | 0.005176 | 5.4 | 6.80E-08 | ***
c
Mean Value of log2FC
Time
P1vsN1
P2vsN2
P3.6vsN3.6
Cluster
1
2
3
4
5
6
7
8
d
Cluster 2 (212 genes)
Stress response: pathogen
Metabolism: lipid
Stress response: detoxification
log2FC
P1vsN1
P2vsN2
P3.6vsN3.6
Cluster 7 (39 genes)
Stress response: detoxification
Metabolism: lipid
log2FC
P1vsN1
P2vsN2
P3.6vsN3.6
Count
150
100
75
50
25
10
5
2
Zero
P value
10-40
10-20
10-10
10-5
0.001
0.05
NS
e
P1vsN1.up
1442
29
18
12
13
14
194
P2vsN2up
P3.6vsN3.6up
f
1326
P1vsN1down
63
21
22
26
16
161
P2vsN2down
P3.6vsN3.6down

**Supplementary Figure 4: No global shift in HS-induced changes in RNA expression and chromatin accessibility between the primed and naive groups in *glp-1(ts)*.**

**(a)** Venn diagrams display the number of significantly differentially expressed genes (top panel) or peaks (bottom panel) between primed and naive groups upon a 3-hour HS (light grey), and their overlap with the 6-hour HS (dark grey) in the indicated comparisons. A Fisher exact test was conducted to determine if the overlaps between the 3-hour and 6-hour HS are statistically significant; p-values are displayed. **(b)** Scatter plots display genes (top panel) or peaks (bottom panel) with significant changes identified in naive or primed groups after a 6-hour HS. The red line indicates the linear regression line with a 95% confidence level, and the black dashed line indicates identity line where x=y. Pearson correlation, R-square values, and coefficient tables are displayed. **(c)** The plot illustrates the temporal dynamics of RNA expression differences between primed and naive worms. The y-axis represents time, including timepoint 1 (P1 vs. N1), timepoint 2 (P2 vs. N2), and timepoint 3 (P3.6 vs. N3.6), while the x-axis represents the mean log2FC. The plot includes significant differentially expressed genes identified at any one of the three timepoints, which were grouped into eight trajectories using K-means clustering analysis. Only the mean log2FC for each trajectory is displayed as a representative trend. Trajectories of all genes in clusters 2 and 7 are displayed in **(d)**. Wormcat GO enrichment analysis for genes in clusters 2 and 7 is also shown. Wormcat p-values are determined by one-sided Fisher test with FDR correction. Gene lists for each cluster in (c–d) are provided in *source.data*. Venn diagrams display the number of significantly upregulated **(e)** or downregulated **(f)** genes identified at timepoint 1 (P1 vs N1) and their overlap with timepoint 2 (P2 vs N2) and timepoint 3 (P3.6 vs N3.6) from RNA-seq analysis (Fig. 3a-b, d)

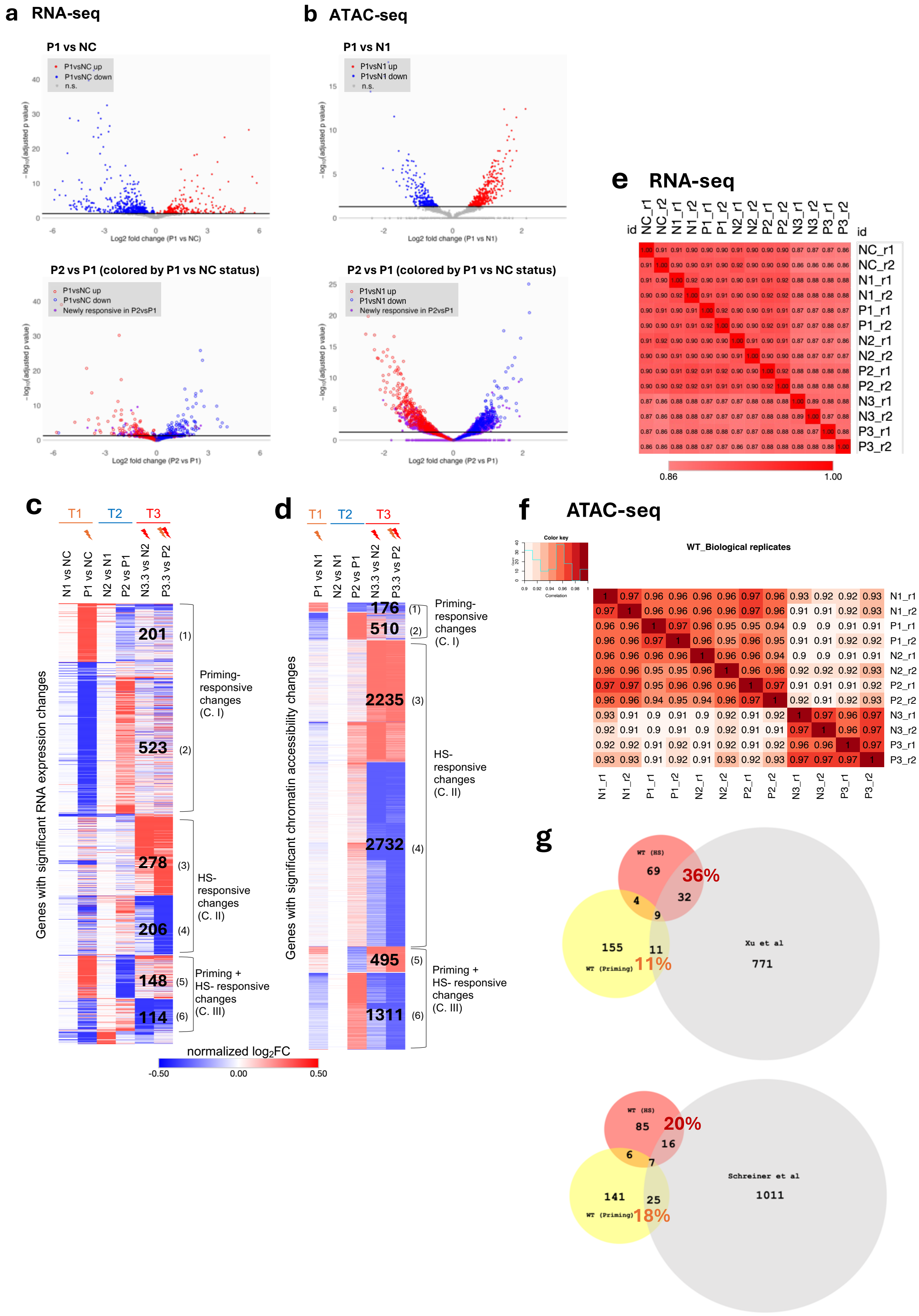
a RNA-seq
P1 vs NC
P1vsNC up
P1vsNC down
n.s.
Log2 fold change (P1 vs NC)
P2 vs P1 (colored by P1 vs NC status)
Newly responsive in P2vsP1
Log2 fold change (P2 vs P1)
b ATAC-seq
P1 vs N1
P1vsN1 up
P1vsN1 down
Log2 fold change (P1 vs N1)
P2 vs P1 (colored by P1 vs NC status)
c
T1
T2
T3
N1 vs NC
P1 vs NC
N2 vs N1
P2 vs P1
N3.3 vs N2
P3.3 vs P2
Genes with significant RNA expression changes
201 (1)
523 (2)
Priming-responsive changes (C. I)
278 (3)
206 (4)
HS-responsive changes (C. II)
148 (5)
114 (6)
Priming + HS- responsive changes (C. III)
normalized log2FC
-0.50
0.00
0.50
d
P1 vs N1
N2 vs N1
P2 vs P1
N3.3 vs N2
P3.3 vs P2
Genes with significant chromatin accessibility changes
176 (1)
510 (2)
Priming-responsive changes (C. I)
2235 (3)
2732 (4)
HS-responsive changes (C. II)
495 (5)
1311 (6)
Priming + HS- responsive changes (C. III)
e RNA-seq
id
NC_r1
NC_r2
N1_r1
N1_r2
P1_r1
P1_r2
N2_r1
N2_r2
P2_r1
P2_r2
N3_r1
N3_r2
P3_r1
P3_r2
0.86
1.00
f ATAC-seq
Color key
Correlation
WT_Biological replicates
N1_r1
N1_r2
P1_r1
P1_r2
N2_r1
N2_r2
P2_r1
P2_r2
N3_r1
N3_r2
P3_r1
P3_r2
g
WT (HS)
69
36%
32
4
9
155
11
WT (Priming)
11%
Xu et al
771
WT (HS)
85
20%
16
6
7
141
25
WT (Priming)
18%
Schreiner et al
1011

**Supplementary Figure 5: Priming-responsive changes largely restore after recovery and are distinct from Heat Shock-responsive changes in gene expression and chromatin accessibility in WT.**

**(a)** RNA-seq and **(b)** ATAC-seq volcano plots. Top panels: Differential gene expression or chromatin accessibility after priming (a: P1 vs NC; b: P1 vs N1). Red and blue indicate significantly up- or downregulated genes/peaks, respectively (adjusted $p < 0.05$, threshold shown as black horizontal line). *Bottom panels:* Differential gene expression or chromatin accessibility after recovery (P2 vs P1), with genes/peaks colored by their P1 vs NC or P1 vs N1 status. Red and blue open circles represent genes/peaks previously up- or downregulated at priming, showing largely opposite regulation after recovery. Purple points indicate "newly responsive" genes/peaks that were not significantly changed in priming but became differentially regulated in P2 vs P1. For RNA-seq **(a)**, the y-axis was truncated for visualization (Max = 45). Because NC timepoints were not collected in ATAC-seq, N1 was used as the baseline comparison. Heatmaps display genes with significant RNA expression changes **(c),** or chromatin accessibility changes **(d)** identified across the indicated comparisons, clustered by K-mean analysis using Morpheus. The colors represent normalized log2FC. The clusters in the heatmaps are arranged to parallelly present shared patterns between changes in RNA expression and chromatin accessibility. Heatmaps are classified into three categories. Category I (C. I), Priming-responsive changes: Involved clusters (1) and (2) in both **(c), (d)**; Category II (C. I), HS-responsive changes: Involved clusters (3) and (4) in both **(c), (d)**; Category III (C. III), Priming + HS-responsive changes: Involved clusters (5) and (6) in both **(c), (d)**. Number indicates the number of genes in the clusters (Details of the gene lists in the heatmaps can be found in the *source.data*). Quality control for transcriptomic and chromatin accessibility profiles of WT*:* **(e)** Spearman's correlation analysis of RNA-seq profiles for all samples, including negative control (NC), naive (N), and primed (P) groups at timepoints 1 to 3 across independent replicates; **(f)** Pearson's correlation analysis of ATAC-seq profiles for all samples, including naive (N) and primed (P) groups at timepoints 1 to 3 across independent replicates. **(g)** Venn diagrams showing overlaps between priming-induced upregulated genes (yellow) and HS-induced upregulated genes (red) identified in WT in this study, and HS-upregulated genes (gray) from published datasets by Xu *et al.* (left) and Schreiner *et al.* (right). Percentages indicate the proportion of shared HS-upregulated genes between studies. Experimental conditions are indicated below: **WT (Priming):** N2, day 1 adults, 30 °C for 6hr vs. 20 °C (P1 vs NC). **WT (HS):** N2, day 1 + 12 h adults, 35 °C for 3hr vs. 20 °C (N3.3 vs N2). **Xu et al.:** N2, day 1 adults, 35 °C for 1 h vs. 20 °C. **Schreiner et al.:** N2, L4 larvae, 35 °C for 4 h vs. 20 °C.

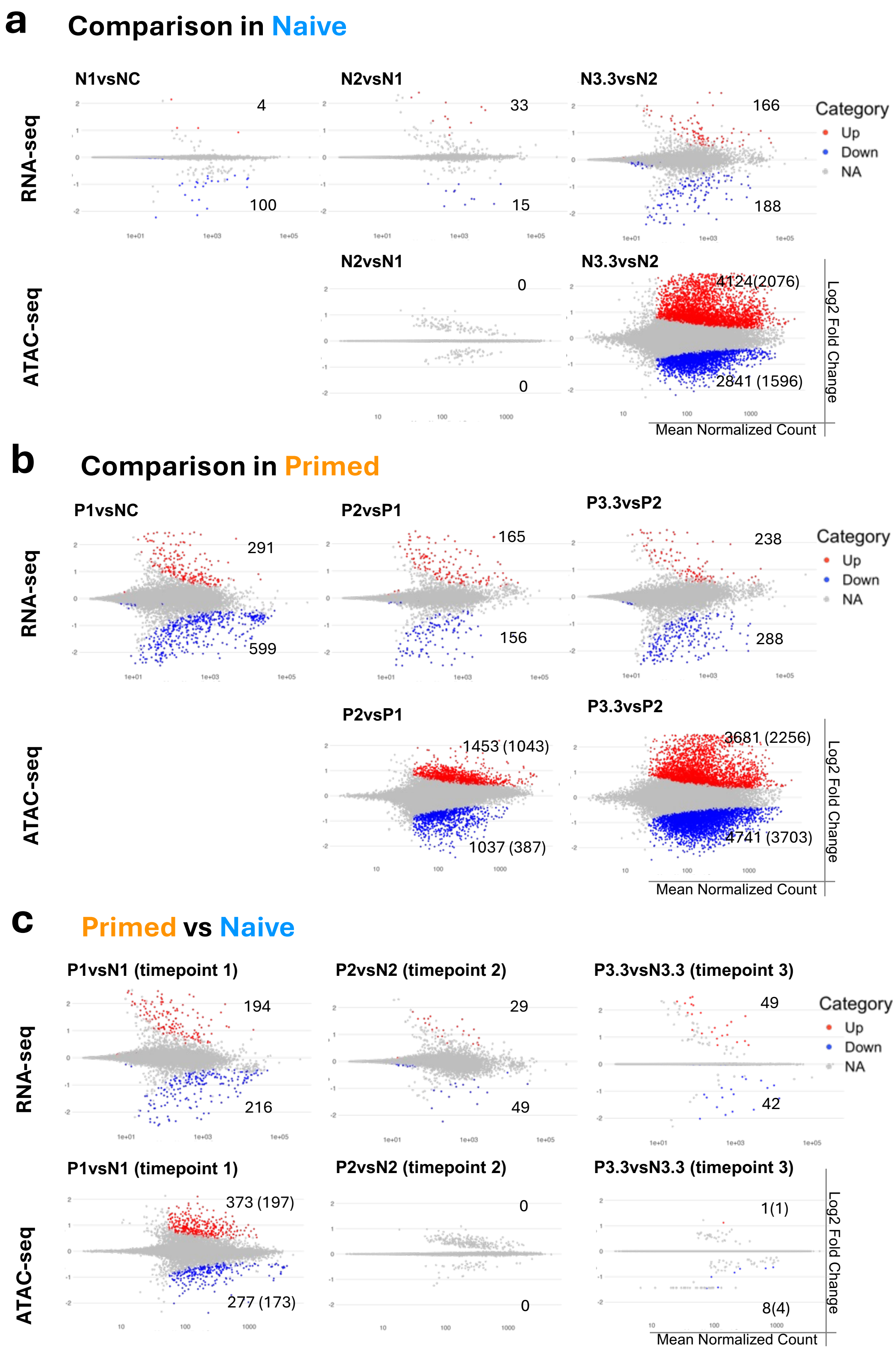
a
Comparison in Naive
N1vsNC
N2vsN1
N3.3vsN2
RNA-seq
4
100
33
15
166
188
Category
Up
Down
NA
ATAC-seq
N2vsN1
N3.3vsN2
0
0
4124(2076)
2841 (1596)
Log2 Fold Change
Mean Normalized Count
b
Comparison in Primed
P1vsNC
P2vsP1
P3.3vsP2
RNA-seq
291
599
165
156
238
288
Category
Up
Down
NA
ATAC-seq
P2vsP1
P3.3vsP2
1453 (1043)
1037 (387)
3681 (2256)
4741 (3703)
Log2 Fold Change
Mean Normalized Count
c
Primed vs Naive
P1vsN1 (timepoint 1)
P2vsN2 (timepoint 2)
P3.3vsN3.3 (timepoint 3)
RNA-seq
194
216
29
49
49
42
Category
Up
Down
NA
ATAC-seq
P1vsN1 (timepoint 1)
P2vsN2 (timepoint 2)
P3.3vsN3.3 (timepoint 3)
373 (197)
277 (173)
0
0
1(1)
8(4)
Log2 Fold Change
Mean Normalized Count

**Supplementary Figure 6: Differential analyses across timepoints and correlation between RNA expression and chromatin accessibility in WT.**

MA plots display log2FC of gene expression from RNA-seq (top panel) and chromatin accessibility from ATAC-seq (bottom panel) for the indicated comparison in naive **(a)** and primed **(b)**, and comparison between two groups **(c)**. Differential analyses were calculated using DESeq2. Significant changes (p-adj < 0.05) are marked in red for upregulation (log2FC > 0) and in blue for downregulation (log2FC < 0), while unchanged are marked in grey (p-adj >= 0.05). Numbers indicate the count of significant genes for RNA-seq data and significant peaks (and their associated genes in brackets) for ATAC-seq data for each plot.

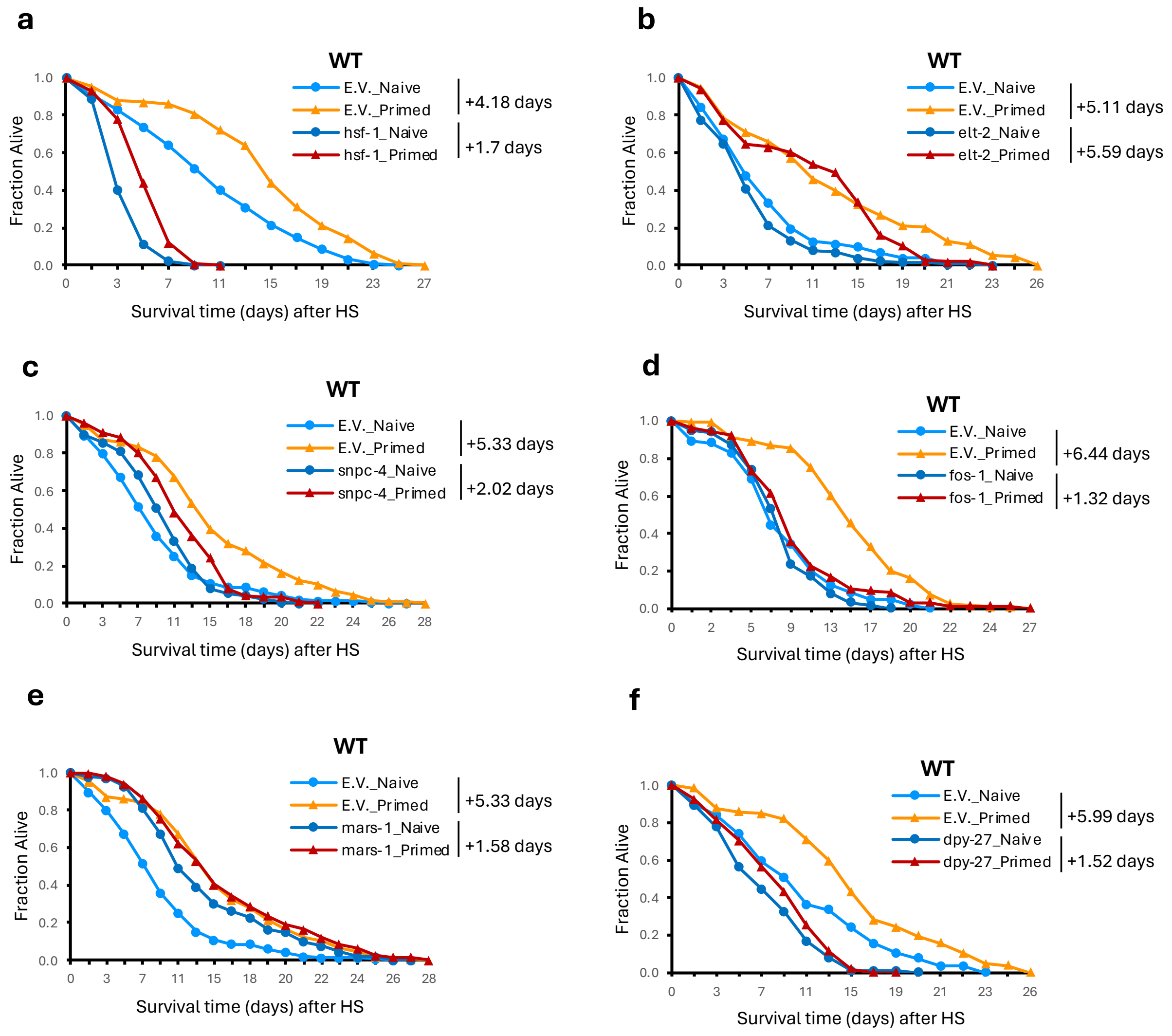


**Supplementary Figure 7: Key regulators of heat hormesis**

Thermal-resistance was assessed based on the survival of WT worms treated with the indicated RNAi and after being subjected to our hormesis regimen and challenged with either 3-hour or 4.5-hour HS. Survival curves represent combined data from multiple independent experiments (N) for *WT*_Naive or *WT*_Primed treated with empty vector (E.V.) control RNAi or *hsf-1* RNAi **(N=3)**, *elt-2* RNAi **(N=2),** *snpc-4* RNAi **(N=3)**, *fos-1* RNAi **(N=4)**, *mars-1* RNAi **(N=3)**, and *dpy-27* RNAi **(N=2).** The mean survival extension (in days) for each condition is indicated. Details in Supplementary Data 5.

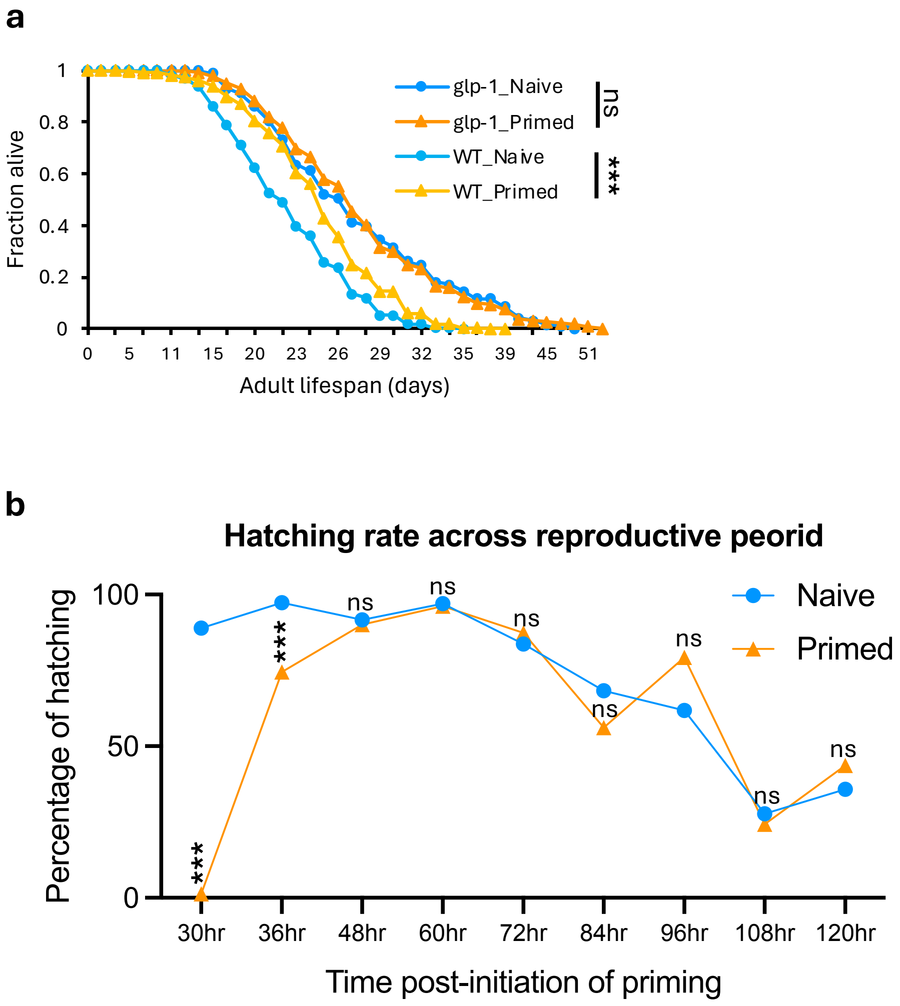


**Supplementary Figure 8. Impact of 30 °C priming on lifespan and reproduction.** **(a)**Lifespan of *glp-1(ts)* or WT worms with or without priming was assessed at 20°C. The figure represents combined data from four independent experiments (N=4). Survival curves for *glp-1*_Naive, *glp-1*_Primed, WT_Naive, WT_Primed are shown. WT worms cultured at 25 °C from eggs to the L4 stage and then shifted to 20 °C overnight prior to priming (to match the *glp-1(ts)* culturing conditions) exhibited a smaller lifespan extension compared to worms continuously cultured at 20 °C (Fig. 5b). **(b)** Hatching rate for each indicated time period was calculated based on the number of eggs laid and the number of hatched larvae 24 hours later. Analyses were performed in WT worms with three biological replicates (N = 3) totaling 25 individuals. Log-rank test was used to compare mean lifespan in (a). Two-tailed unequal variances t-tests were used in (b). *** indicates p < 0.001. Details are provided in source.data.

#### Descriptions of additional Supplementary files

**File Name: Supplementary Data 1**

**Description:** Gene lists and log2FC values for the indicated comparisons of K-means clusters, originally identified from Morpheus (https://software.broadinstitute.org/morpheus) and further rearranged into a heatmap corresponding to Figure 2c.

**File Name: Supplementary Data 2**

**Description:** Peak (associated-gene) lists and log2FC values for the indicated comparisons of K-means clusters, originally identified from Morpheus (https://software.broadinstitute.org/morpheus) and further rearranged into a heatmap corresponding to Figure 2d.

**File Name: Supplementary Data 3**

**Description:** Gene ontology (GO) analysis summary for the indicated gene lists, identified using WormCat 2.0 (Category 2 and 3)(5), including statistical details, corresponding to Figure 2e-g and Figure 3e-f.

**File Name: Supplementary Data 4**

**Description:** RNAi screening list, including instructions for RNAi feeding time and raw data of individual RNAi gene KD survival results.

**File Name: Supplementary Data 5**

**Description:** RNAi screening list in WT background, including instructions for RNAi feeding time and raw data of individual RNAi gene KD survival results.

**File Name: Supplementary Data 6**

**Description:** ATAC-seq analysis pipeline and istructions for execution in Linux

**File Name: Supplementary Data 7**

**Description:** RNA-seq analysis pipeline and instructions for execution in Linux

**File Name: Supplementary Data 8**

**Description:** R script for ruunning differential expression analysis and generating an MA and Volcano plot.

**File Name: Supplementary Data 9**

**Description:** R script for annotating consensus peaks.

**File Name: Supplementary Data 10**

**Description:** R script for fetching promoter region sequences for motif analysis.

**File Name: Supplementary Data 11**

**Description:** R script for conducting temporal dynamic analysis with time-series clustering and plotting interactive trajectories.

**File Name: Supplementary Data 12**

**Description:** Matrix of raw read counts from RNA-seq data. (Will also be uploaded to GEO metadata.)

**File Name: Supplementary Data 13**

**Description:** Matrix of raw read counts in identified consensus peaks from ATAC-seq data. (Will also be uploaded to GEO metadata.)

**File Name: Supplementary Data 14**

**Description:** Reference Elements for peak annotation, adapted from a published dataset(2).

#### References


1. Yan F, Powell DR, Curtis DJ, Wong NC. From reads to insight: a hitchhiker's guide to ATAC-seq data analysis. Genome Biol [Internet]. 2020 Feb 3 [cited 2025 Feb 16];21(1):22. Available from: https://doi.org/10.1186/s13059-020-1929-3
2. Chen RAJ, Down TA, Stempor P, Chen QB, Egelhofer TA, Hillier LW, et al. The landscape of RNA polymerase II transcription initiation in C. elegans reveals promoter and enhancer architectures. Genome Res. 2013 Aug;23(8):1339–47.
3. Daugherty AC, Yeo RW, Buenrostro JD, Greenleaf WJ, Kundaje A, Brunet A. Chromatin accessibility dynamics reveal novel functional enhancers in C. elegans. Genome Res. 2017 Dec;27(12):2096–107.

4. Jänes J, Dong Y, Schoof M, Serizay J, Appert A, Cerrato C, et al. Chromatin accessibility dynamics across C. elegans development and ageing. Lee SS, Tyler JK, editors. eLife [Internet]. 2018 Oct 26 [cited 2025 Feb 16];7:e37344. Available from: https://doi.org/10.7554/eLife.37344

5. Holdorf AD, Higgins DP, Hart AC, Boag PR, Pazour GJ, Walhout AJM, et al. WormCat: An Online Tool for Annotation and Visualization of Caenorhabditis elegans Genome-Scale Data. Genetics [Internet]. 2020 Feb 1 [cited 2025 Feb 16];214(2):279–94. Available from: https://doi.org/10.1534/genetics.119.302919
